# DNMT mRNA stability and YB-1 cooperatively regulate ABCB1 to drive cisplatin chemoresistance in cholangiocarcinoma

**DOI:** 10.64898/2026.06.05.730343

**Authors:** Tao Lin, Chenjun Huang, Chenhao Tong, Heng Liu, Roman Liebe, Lungen Lu, Jun Li, Jonathan A. Lindquist, Matthias PA Ebert, Peter R Mertens, Steven Dooley, Hong-Lei Weng

## Abstract

**Background and Aims:** Intrahepatic cholangiocarcinoma (iCCA) is a tumor type with a high lethality due to late diagnosis and profound resistance to conventional chemotherapy. To date the molecular mechanisms underlying multidrug resistance remain poorly defined. Here, we integrate single-cell transcriptomics, clinicopathological analysis, and functional genomics to elucidate the molecular basis of cisplatin resistance in iCCA.

**Approach and Results:** Single-cell RNA sequencing of iCCA and adjacent liver tissues revealed pronounced expression of Y-box–binding protein 1 (YB-1) in aneuploid malignant cholangiocarcinoma cells, with YB-1 expression progressively increasing during malignant evolution and strongly associated with chemoresistance. Clinically, elevated YB-1 expression-particularly its nuclear localization-robustly predicts poor overall survival and chemotherapy failure in patients with iCCA. Mechanistically, we demonstrate that cisplatin induces phosphorylation-dependent nuclear translocation of YB-1, enabling direct transcriptional activation of the drug efflux transporter ABCB1. Importantly, this process requires ABCB1 promoter demethylation, which is driven by cisplatin-induced, m⁶A-dependent destabilization of DNMT1 and DNMT3B mRNAs. This destabilization occurs through disruption of the YB-1-IGF2BP1/3-DNMT mRNA stabilizing complex and subsequent recruitment of DNMT transcripts to YTHDF2-mediated processing bodies for degradation.

**Conclusions:** Our findings uncover a previously unrecognized YB-1-m⁶A-DNMT regulatory axis that drives chemotherapeutic resistance in iCCA, highlighting YB-1 as both a prognostic biomarker and a promising therapeutic target.

## Introduction

Intrahepatic cholangiocarcinoma (iCCA) is the second most common primary malignant liver tumor, posing a heavy burden on global health due to its rising incidence and extremely poor prognosis^1–3^. Characterized by high genetic heterogeneity and dense, pro-fibrotic stroma, iCCA is often diagnosed at an advanced stage, making surgical resection, the only possible cure, infeasible for most patients^2,4,5^. For these patients, systemic chemotherapy, such as gemcitabine- and cisplatin-based regimens, remains the standard of care^6,7^. However, the clinical efficacy of these chemotherapies is severely limited by intrinsic and acquired multidrug resistance, resulting in a five-year survival rate of less than 10%^8–11^. Genetic heterogeneity is one of the major causes of drug resistance^7^. Despite extensive genomic analyses, the precise molecular mechanisms controlling gemcitabine and cisplatin resistance remain elusive.

Y-box-binding protein 1 (YB-1) plays a crucial role in cancer progression and stress response^12–14^. As a multifunctional cold shock protein, it functions both as a transcription factor in the nucleus and as a translation regulator in the cytoplasm^14–16^. YB-1 is known to be overexpressed in various solid tumors and participates in promoting epithelial-mesenchymal transition (EMT), cell proliferation, and DNA repair^12,13,17–19^. More importantly, YB-1 is associated with the activation of the ABCB1 gene, which encodes a membrane-bound efflux pump multidrug resistance protein 1 (MDR1), that actively expels chemotherapeutic drugs from cells^20–23^. In addition to transcription factor-mediated regulation, ABCB1 expression is also controlled by epigenetic mechanisms, including promoter DNA methylation, which can influence its transcriptional activity^23–25^. Although the association between YB-1 and drug resistance has been confirmed in various cancers, how it regulates ABCB1 gene remains uncertain^26,27^.

N6-methyladenosine (m^6^A) has been identified as a key post-transcriptional regulator of mRNA stability^28,29^. m^6^A modification, regulated by "writers" (e.g., METTL3/14), "erasers" (e.g., FTO), and "readers" (e.g., YTHDF2, IGF2BPs), has become a key regulator of oncogene and tumor suppressor gene expression^30–32^. However, the interaction between the m^6^A signaling pathway and DNA methylation to regulate the YB-1-driven resistance program in iCCA remains unexplored.

In this study, we employed a comprehensive approach combining single-cell RNA sequencing, clinical cohort analysis, and functional genomics to elucidate the drivers of cisplatin resistance in iCCA. YB-1 was identified as a biomarker for malignant aneuploid cholangiocarcinoma cells. Cisplatin-induced phosphorylation-dependent nuclear translocation of YB-1 is crucial for ABCB1 transcription. Furthermore, cisplatin induces m^6^A-dependent degradation of DNA methyltransferases DNMT1 and DNMT3B, thereby promoting ABCB1 promoter demethylation and enhancing YB-1-mediated multidrug resistance transcriptional activation.

## Results

### Single-cell transcriptome analysis reveals upregulation of YB-1 in cholangiocarcinoma cells

To characterize the cellular composition and molecular features of cholangiocarcinoma, we analyzed publicly available single-cell transcriptome data from intrahepatic cholangiocarcinoma (iCCA) and adjacent non-tumor liver tissue (GSE138709^33^). UMAP^34^ visualization of the integrated scRNA-seq dataset, comprising 33,990 cells from eight samples (five tumor and three adjacent; 17,089 tumor cells vs. 16,901 adjacent cells), identified 21 distinct clusters (**Figure S1A**). Cell type annotation based on classical marker gene expression revealed 13 major cell lineages, including epithelial, endothelial, stromal, and immune cells (**Figure 1A**, **Figure S1B**). These populations exhibited distinct spatial organization in UMAP space, reflecting their unique transcriptomic profiles. For example, cholangiocytes specifically expressed SOX9, KRT19, and EPCAM, whereas hepatocytes expressed ALB, CYP3A4, and APOA1 (**Figure S1B**). B cells expressed MS4A1, CD19, CD79A, and CD79 (**Figure S1B**).

**Figure 1.**
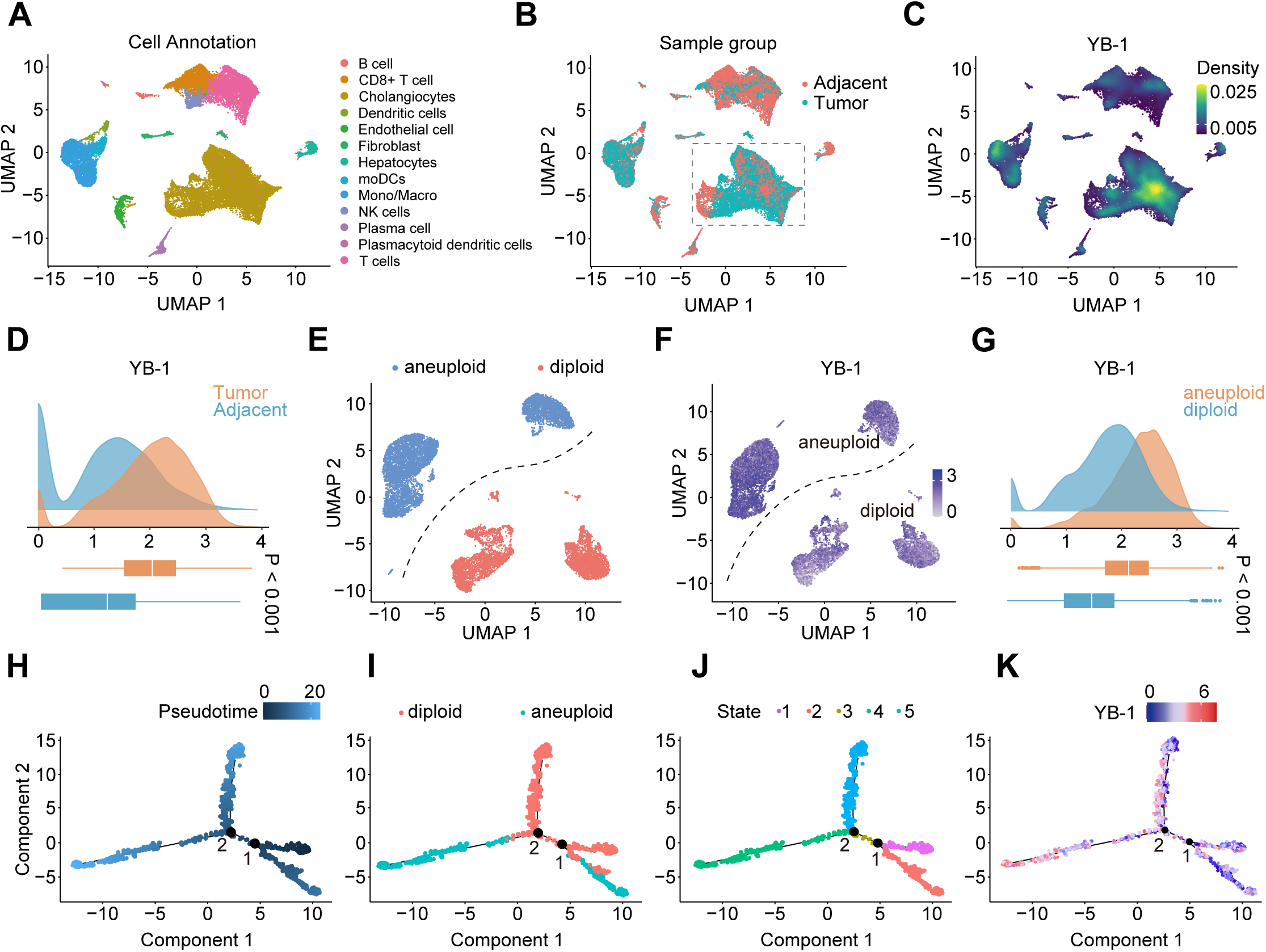
Single-cell landscape of YB-1 expression, copy number variation, and malignant cell trajectories in tumor and adjacent tissues. **(A)** Cell type annotation of the UMAP plot reveals major cellular populations, including immune cells (B cells, CD8+ T cells, T cells, NK cells, moDCs, Mono/Macro, plasma cells, dendritic cells), stromal cells (fibroblasts, endothelial cells), and epithelial cells (cholangiocytes, hepatocytes). **(B)** UMAP plot colored by sample origin, distinguishing between tumor (red) and adjacent non-tumor tissue (teal). **(C)** The UMAP density map shows the spatial distribution and expression levels of YB-1. **(D)** Density plots and box plots compared the YB-1 expression in tumor tissues (orange) and adjacent non-tumor tissues (blue) (p < 0.0001). **(E)** UMAP plot showing inferred copy number variation status in cholangiocytes, distinguishing between diploid (blue) and aneuploid (red) cells. **(F)** UMAP visualization of single-cell transcriptomic data showing YB-1 expression distribution across diploid and aneuploid cells. **(G)** Density plots and box plots compared the YB-1 expression in aneuploid (orange) and diploid (blue) cells (p < 0.0001). **(H)** Pseudo-temporal trajectory analysis of malignant cells rooted in diploid cells. Trajectories are colored according to pseudo-temporal progression (light blue = late pseudo-time, dark blue = early pseudo-time). **(I)** Visualization of pseudo-temporal trajectories colored by copy number status, distinguishing between diploid (red) and aneuploid (cyan) cell populations. **(J)** Unsupervised clustering along the trajectories identifies discrete cell states (states 1-5) and labels the pseudo-temporal trajectories accordingly. **(K)** Pseudo-temporal trajectories superimposed with YB-1 gene expression levels. The color gradient (blue to red) represents the expression intensity of YB-1.

Comparison between tumor tissue and adjacent tissue samples revealed significant differences in cellular composition (**Figure 1B**, **Figure S1C**). Adjacent samples retained a higher proportion of normal cholangiocytes and immune cells, whereas tumor tissues exhibited increased infiltration of immune cells, including B cells, T cells, macrophages, and NK cells (**Figure S1C**). Notably, UMAP density mapping revealed high YB-1 expression within the cholangiocyte cluster (**Figure 1C**). Quantitative analysis and distribution plots further demonstrated that tumor samples exhibited significantly elevated YB-1 expression compared with adjacent tissue (**Figure 1D**, P<0.001).

Subsequently, we performed a comprehensive subclustering analysis to characterize the heterogeneity of malignant cholangiocarcinoma cells. Fourteen distinct subclusters (clusters 0-13) were identified within the cholangiocyte compartments (**Figure S1D**). Cell type annotation further classified these populations into three major categories: cholangiocytes (red), Liver progenitor cells (LPCs, green), and malignant cells (blue) (**Figure S1E**). Cholangiocytes and malignant cells formed well-separated clusters, whereas LPCs occupied a smaller, intermediate cluster (**Figure S1E**). Both cholangiocytes and malignant cells showed strong expression of bile duct epithelial markers SOX9 and KRT19 (**Figure S1F**). Compared with cholangiocytes, malignant cells uniquely expressed oncogenes MYC, BIRC5, UBE2C, CDK1, CCNB1, and CCNB2 (**Figure S1F**). LPCs displayed distinct molecular signatures, including high expression of albumin (ALB) and hepatocyte nuclear factor 4 alpha (HNF4A), consistent with a progenitor cell state with hepatocyte differentiation potential (**Figure S1F**).

To cross-validate this classification, chromosomal copy-number variation (CNVs) were inferred from expression data using inferCNV^35^ to estimate ploidy status. Significant genomic instability was observed in the malignant cell population (**Figure S2**). Reference diploid cells exhibited a uniform expression pattern across chromosomes, with minimal copy-number variation in cholangiocytes (**Figure S2**). In contrast, cholangiocarcinoma cells showed widespread chromosomal aberrations, including extensive copy-number gains (red) and deletions (blue) across multiple regions (**Figure S2**). Ploidy analysis revealed a clear separation between diploid and aneuploid populations in UMAP space (**Figure 1E**). This spatial concordance between transcriptional state, cell identity, and chromosomal stability suggests that aneuploidy is a key molecular feature of the malignant phenotype, distinguishing malignant cholangiocarcinoma from normal cholangiocytes (**Figure 1E**, **Figure S1E**).

Intriguingly, UMAP projection revealed that YB-1 expression was primarily elevated in aneuploid malignant cell clusters, whereas significantly lower expression was observed in diploid populations (**Figure 1F**). Raincloud plot analysis further demonstrated significantly higher YB-1 expression in aneuploid cells than in diploid cells (**Figure 1G**, P<0.001).

To elucidate the dynamic YB-1 expression during cholangiocarcinoma development, we performed pseudo-chronological trajectory analysis (**Figure 1H**). This analysis revealed multiple developmental pathways. Chromosomal profiling and cell state annotation along the trajectory indicated that a subset of diploid cells transitioned into aneuploid populations, whereas others retained diploid characteristics (**Figure 1I-J**). Further trajectory analysis showed relatively low YB-1 expression in the initial diploid cholangiocarcinoma population (**Figure 1K**). During tumor progression, YB-1 expression gradually increased, with a pronounced elevation in aneuploid cells corresponding to later stages of the trajectory (**Figure 1K**). Although YB-1 expression exhibited heterogeneity (**Figure 1K**), its overall levels progressively increase during cholangiocarcinoma development and progression.

### YB-1 expression and nuclear localization are associated with the prognosis of iCCA patients

Next, we analyzed public transcriptome datasets (GSE76311, TCGA-CHOL, and GSE107943). These datasets consistently showed significantly elevated YB-1 expression in tumor samples (p < 0.0001; **Figures 2A–C**). Clinical outcome analysis based on the GSE107943 cohort further revealed that patients with high YB-1 expression had significantly lower overall survival compared with those with low YB-1 expression, with median survival times of approximately 15 months and 60 months, respectively (p=0.0461, Figure 2D).

**Figure 2.**
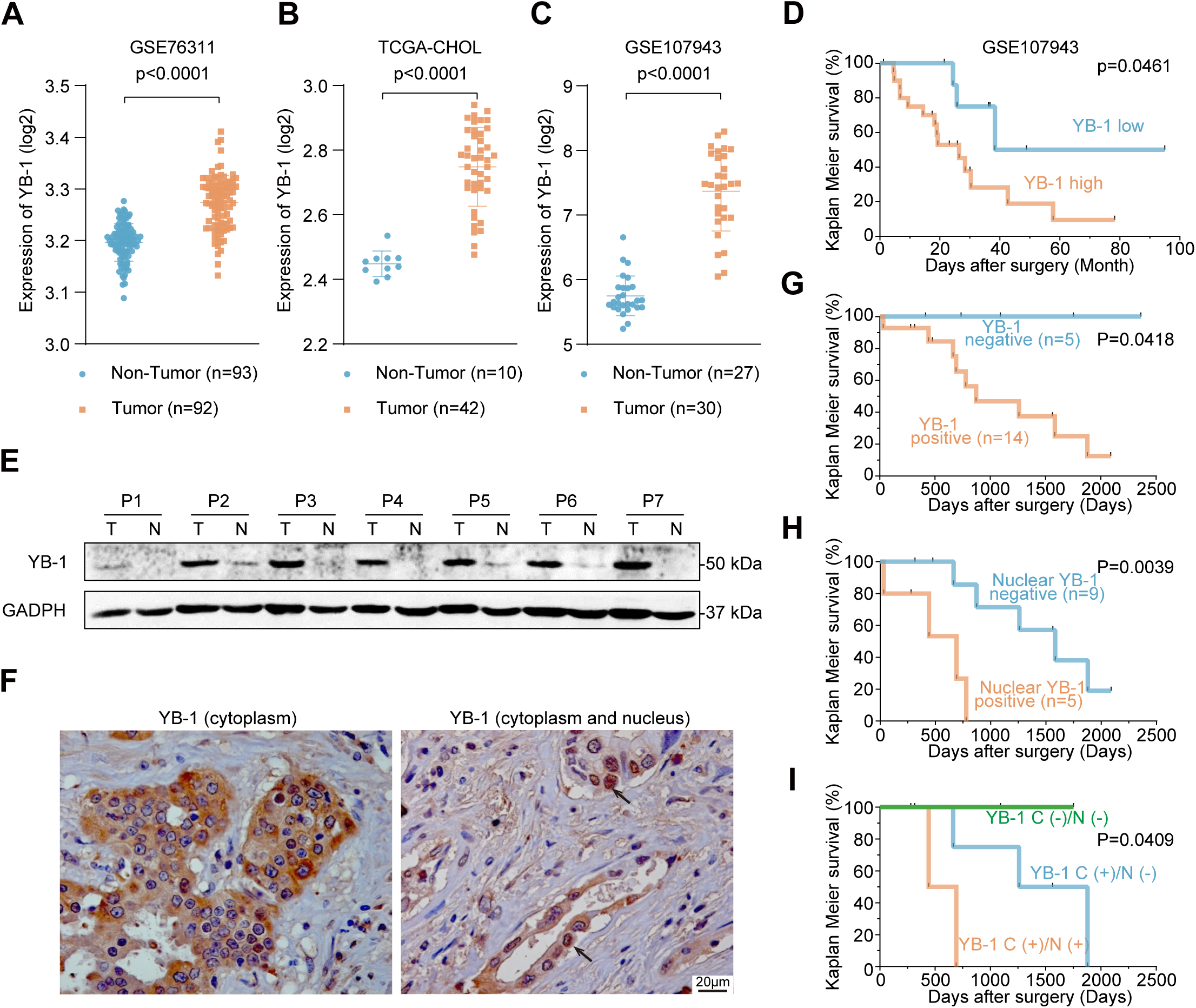
YB-1 expression and localization predict patient survival. **(A-C)** Comparison of YB-1 mRNA expression in tumor samples and non-tumor tissues in transcriptome datasets (GSE76311, TCGA-CHOL, and GSE107943) (all p < 0.0001). **(D)** The Kaplan-Meier survival analysis of the GSE107943 cohort showed the correlation between YB-1 expression and overall survival in patients with cholangiocarcinoma (p=0.0461). **(E)** Western blot analysis of YB-1 protein expression in paired tumor (T) and adjacent non-tumor (N) tissue samples from 7 patients with cholangiocarcinoma (P1-P7). GAPDH serves as a loading control. **(F)** Representative immunohistochemistry images showing YB-1 protein localization in cholangiocarcinoma tissue. **(G)** The Kaplan-Meier survival analysis based on YB-1 expression stratification in immunohistochemistry (Figure 2H) showed the correlation between YB-1 expression and overall survival in patients with cholangiocarcinoma (p=0.0418). **(H)** Kaplan-Meier analysis based on YB-1 subcellular localization showed the relationship between YB-1 expression localization and patient survival (p=0.0039). **(I)** Combined analysis of YB-1 expression and localization showed their correlation with patient survival after chemotherapy (p = 0.0409).

We next performed Western blot analysis to examine YB-1 protein expression in paired liver tissues from seven patients with iCCA. Consistent with the public dataset findings, tumor tissues showed markedly increased YB-1 protein expression compared with adjacent non-tumor tissues (**Figure 2E**). We then performed immunohistochemical staining (IHC) on liver tissue samples from 28 iCCA patients.

Strong YB-1 expression was observed in 21 patients (75%) (**Table 1**). Among these YB-1-positive patients, 13 exhibited YB-1 immunoreactivity exclusively in the cytoplasm of cholangiocarcinoma cells, whereas 8 patients (38.1%) showed YB-1 positivity in both the cytoplasm and nuclei (**Figure 2F** and **Table 1**). Correlation analysis demonstrated that nuclear YB-1 expression in tumor cells was positively associated with tumor invasion (T stage, P = 0.044), vascular invasion (P = 0.015), postoperative metastasis (P = 0.002), and advanced disease stage (P = 0.038). In contrast, cytoplasmic YB-1 expression was associated only with a high incidence of cholestasis (P = 0.040) (**Table 1**).

**Table 1.** Relationships between YB-1 expression and various clinicopathologic factors in ICC patients.

| Clinicopathologic factor | YB-1 expression in cytoplasm |  |  | YB-1 expression in nucleus |  |  |
| --- | --- | --- | --- | --- | --- | --- |
|  | IHC score > 1<br>(n=21) | IHC score ≤ 1<br>(n=7) | *P* | Positive<br>(n=8) | Negative<br>(n=20) | *P* |
| Age (y, mean ± SD) | 60.38 ± 8.65 | 58.43 ± 9.74 | 0.620 | 58.25 ± 11.55 | 60.55 ± 7.68 | 0.542 |
| Gender |  |  |  |  |  |  |
| Male | 9 | 6 | 0.084 | 4 | 11 | 1.000 |
| Female | 12 | 1 |  | 4 | 9 |  |
| TNM stages (AJCC/UICC) |  |  |  |  |  |  |
| pT |  |  |  |  |  |  |
| pT1-2 | 11 | 5 | 0.662 | 2 | 14 | 0.044 |
| pT3-4 | 10 | 2 |  | 6 | 6 |  |
| pN |  |  |  |  |  |  |
| (−) | 14 | 5 | 1.000 | 4 | 16 | 0.328 |
| (+) | 6 | 2 |  | 3 | 4 |  |
| pNx | 2 | 0 |  | 2 | 0 |  |
| pM |  |  |  |  |  |  |
| (−) | 14 | 7 | 0.141 | 4 | 17 | 0.142 |
| (+) | 7 | 0 |  | 4 | 3 |  |
| Degree of differentiation (G) |  |  |  |  |  |  |
| G1 | 2 | 0 | 0.252 | 1 | 1 | 0.558 |
| G2 | 8 | 6 |  | 3 | 11 |  |
| G3 | 6 | 1 |  | 1 | 6 |  |
| Gx | 5 | 0 |  | 3 | 2 |  |
| Lymph node metastasis (L) |  |  |  |  |  |  |
| (−) | 16 | 5 | 0.633 | 6 | 15 | 1.000 |
| (+) | 4 | 2 |  | 2 | 4 |  |
| Lx | 1 | 0 |  | 0 | 1 |  |
| Vascular invasion (V) |  |  |  |  |  |  |
| (−) | 16 | 6 | 0.554 | 4 | 18 | 0.015 |
| (+) | 3 | 0 |  | 3 | 0 |  |
| Vx | 2 | 1 |  | 1 | 2 |  |
| Cholestasis (C) |  |  |  |  |  |  |
| (−) | 16 | 3 | 0.040 | 7 | 12 | 0.273 |
| (+) | 1 | 3 |  | 0 | 4 |  |
| Cx | 4 | 1 |  | 1 | 4 |  |
| Satellite lesions (S) |  |  |  |  |  |  |
| (−) | 11 | 6 | 0.129 | 3 | 14 | 0.156 |
| (+) | 8 | 0 |  | 4 | 4 |  |
| Sx | 2 | 1 |  | 1 | 2 |  |
| Postoperative Metastasis |  |  |  |  |  |  |
| (−) | 11 | 3 | 0.678 | 0 | 14 | 0.002 |
| (+) | 9 | 4 |  | 7 | 6 |  |
| Mx | 1 | 0 |  | 1 | 0 |  |
| Clinical stages |  |  |  |  |  |  |
| I+II | 8 | 5 | 0.198 | 1 | 12 | 0.038 |
| III+IV | 13 | 2 |  | 7 | 8 |  |

Abbreviations: pTx, pNx, pMx, Gx, Lx, Vx, Cx, Fx and Mx meant data missing cases

Among 28 iCCA patients, 19 (14 with YB-1-positive tumor cells and 5 without YB-1 expression) were followed for six years. At the end of follow-up, all five patients without YB-1-expression were alive (**Figure 2G**). In contrast, among the 14 patients with YB-1 expression, only 5 (35.7%) were survival (**Figure 2G**). We further stratified these 14 patients based on nuclear YB-1 immunopositivity (nuclear-positive vs. nuclear-negative: 5 vs. 9). Survival analysis showed that patients with nuclear YB-1 expression had significantly shorter survival compared with those without nuclear expression (**Figure 2H**, p<0.001). Among the 9 patients without nuclear YB-1 expression, 8 had a survival time longer than 2 years, and 2 remained alive at the end of follow-up (> 5 years) (**Figure 2H**). In contrast, among the 5 patients with nuclear YB-1 expression, 4 died within 2 years and 1 died within 3 years after surgery (**Figure 2H**).

In this cohort, 10 patients received postoperative chemotherapy (gemcitabine and/or cisplatin). Among them, 2 tumors lacked YB-1 expression, 5 showed cytoplasmic YB-1 expression, and 3 exhibited nuclear YB-1 expression (**Table 2**). At the end of follow-up, 2 patients without YB-1 expression were alive, whereas only 2 out of 5 patients (40%) with cytoplasmic YB-1 expression and 1 of 3 patient (33.3%) with nuclear YB-1 expression survived (**Table 2**). Notably, patients with nuclear YB-1 expression had the shortest survival time (**Figure 2I**, p < 0.05). These results suggest that YB-1 expression and its subcellular localization may contribute to cancer progression and chemotherapy resistance.

**Table 2.**
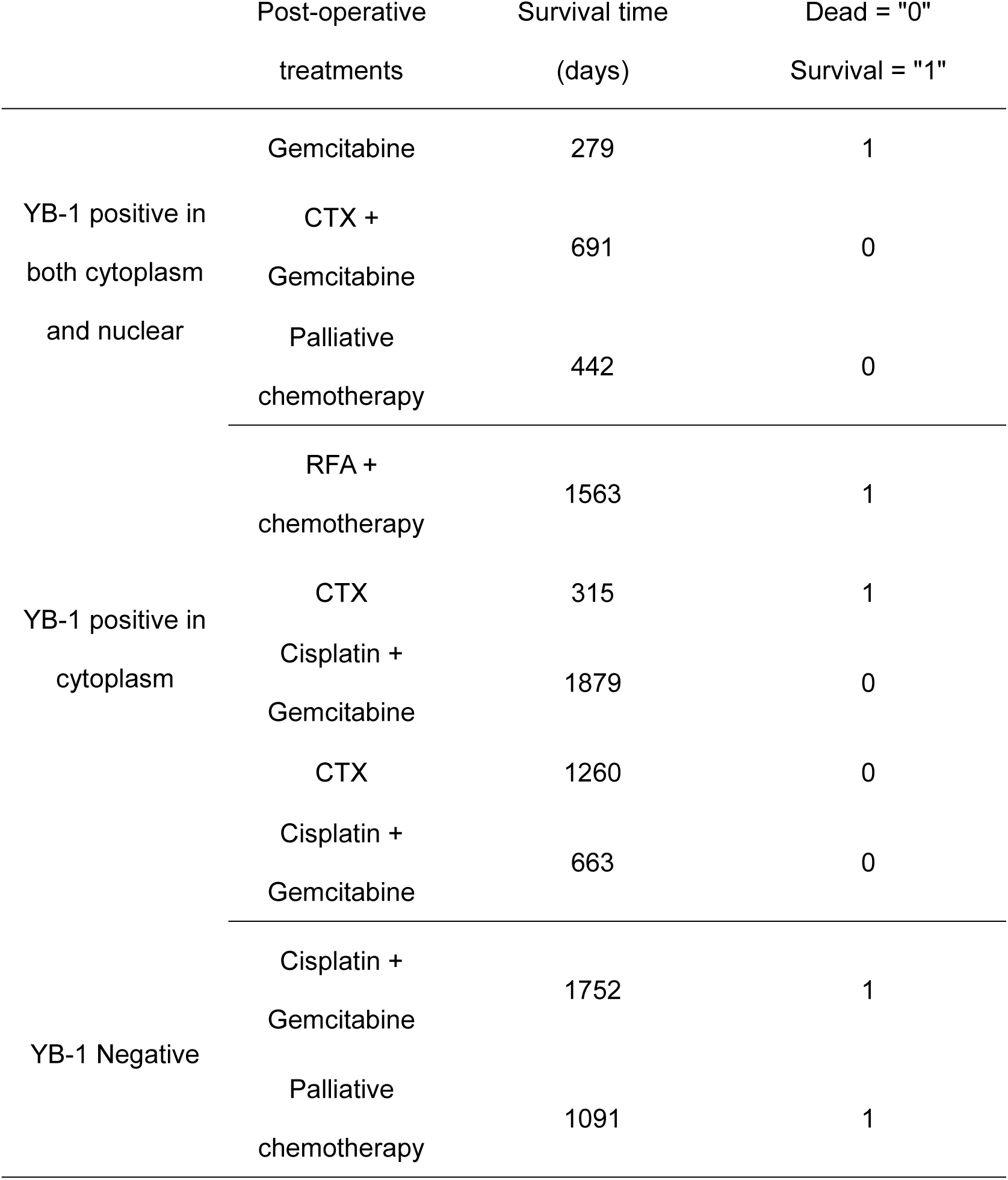
Outcome of CCA patients who received chemotherapy after operation.

### Activation of the YB-1 regulatory network drives multidrug chemotherapy resistance in cholangiocarcinoma cells

To elucidate the molecular mechanisms underlying YB-1-mediated chemotherapy resistance, we quantified regulon activity using SCENIC and derived per-cell regulon activity scores with AUCell^36^. Heatmap analysis revealed differential expression of YB-1 and other transcription factors between aneuploid and diploid cell populations (**Figure 3A**). Notably, both YB-1 expression and YB-1 regulon activity were significantly higher in the aneuploid population than in the diploid population (**Figure 3A-B**, P < 0.0001). Consistently, GSVA scoring also showed elevated YB-1 relevant pathway activity in aneuploid cells (**Figure 3C**, P < 0.0001).

**Figure 3.**
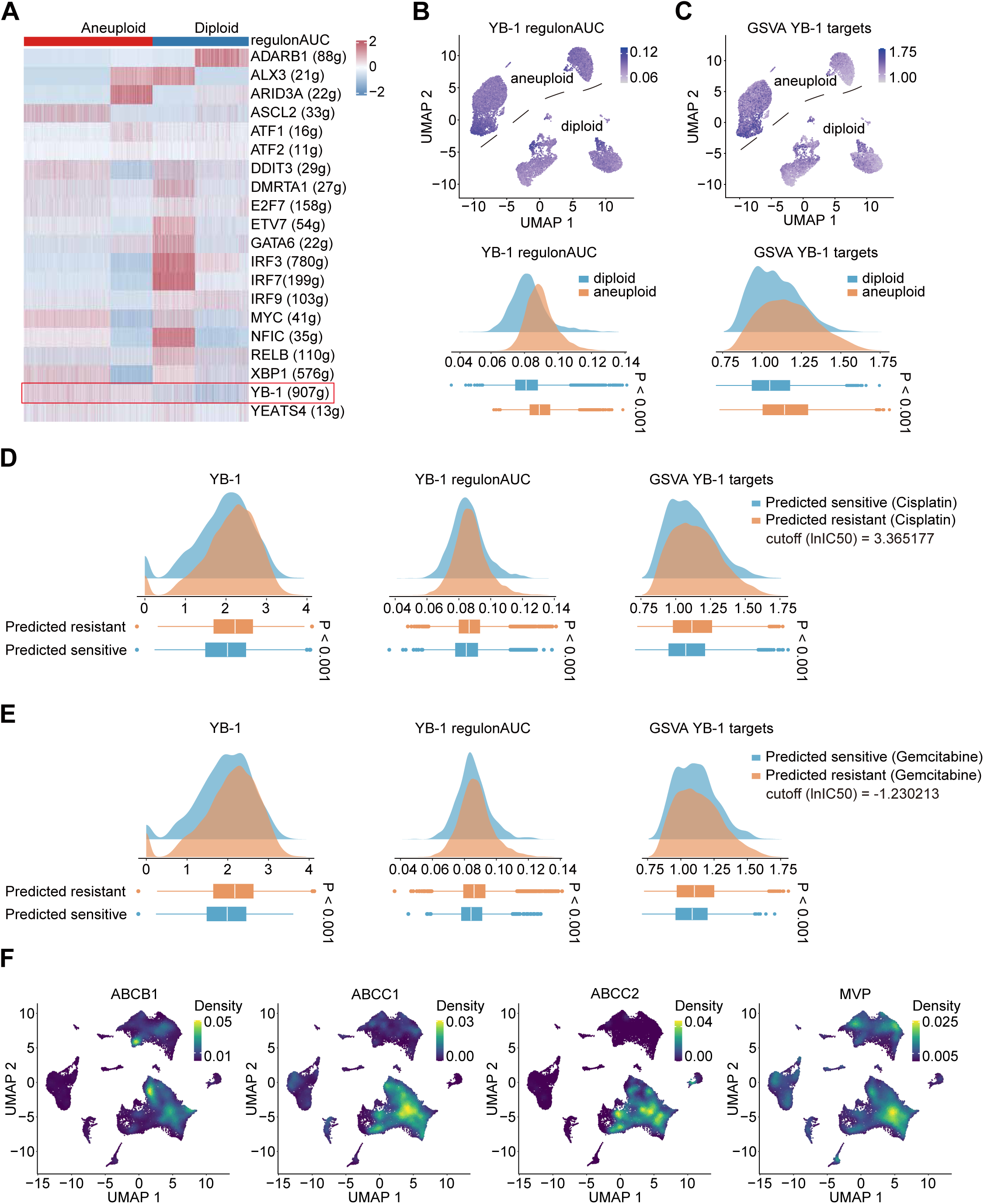
Enhanced YB-1 regulatory activity predicts chemotherapy sensitivity. **(A)** Heatmap displaying the differential expression of transcription factor regulon activities between aneuploid and diploid cells. Each row represents a specific transcription factor regulon with the number of target genes indicated in parentheses. The color scale (blue to red) represents regulon activity scores. **(B)** Top: UMAP visualization showing YB-1 regulon activity (YB-1_regulonAUC) across aneuploid and diploid cells. Bottom: Density plot and box plot comparing YB-1 regulon AUC scores between diploid and aneuploid cells (P < 0.001). **(C)** Top: UMAP visualization displaying the activity score of YB-1 target genes (GSVA_YB-1_targets) in aneuploid and diploid cells. Bottom: Density plot and box plot comparing YB-1 target gene expression between diploid and aneuploid cells (P < 0.001). **(D)** Prediction of cisplatin sensitivity based on YB-1-related features. Density plots and box plots show the distribution of YB-1 expression (left), YB-1 regulon AUC (middle), and GSVA YB-1 target gene scores (right) for cells predicted to be cisplatin-sensitive (blue) versus cisplatin-resistant (orange). The InferCNV cutoff for prediction is 3.365177. **(E)** Prediction of gemcitabine sensitivity based on YB-1-related features. Similar to panel D, the three metrics (YB-1 expression, regulon AUC, and target gene scores) are displayed for predicted gemcitabine-sensitive (blue) versus resistant (orange) cells. The InferCNV cutoff is -1.230213. **(F)** The UMAP density map shows the spatial distribution and expression levels of chemotherapy resistance-related transporter genes. The color gradient (purple to yellow) represents cell density and gene expression levels. ABCB1 (MDR1/P-glycoprotein), ABCC1 (MRP1), ABCC2 (MRP2), and MVP (major vesicle protein).

Next, we used oncoPredict to assess the relationship between YB-1 expression and therapeutic response to gemcitabine or cisplatin, as well as to explore its potential regulatory network associated with chemotherapy response. Based on oncoPredict-estimated IC_50_ values^37^, cancer cells were categorized as predicted sensitive or resistant to gemcitabine or cisplatin. Volcano plot analysis revealed significantly different transcription factor regulons between cisplatin-predicted sensitive and resistant groups (**Figure S3A**). YB-1-associated transcription factor regulons were enriched in the cisplatin resistant group (**Figure S3A**). The regulon AUC heatmap also indicated higher YB-1 regulon activity in predicted resistant cells (**Figure S3B**).

Analyses based on predicted cisplatin IC_50_ values and YB-1 regulon activity showed similar spatial distributions in UMAP space, with greater enrichment of YB-1 activity in aneuploid cells (**Figure S3C-D**). Predicted resistant cells exhibited higher YB-1 expression, regulon activity, and target gene set scores than predicted sensitive cells (**Figure 3D**, all P < 0.0001). A similar pattern was observed in gemcitabine response prediction: resistant cells exhibited higher YB-1 expression, enhanced regulator activity, and increased target gene scores (**Figure 3E**). These results further indicate that activation of the YB-1 regulatory network may play a critical role in chemotherapy resistance.

To identify potential effectors of YB-1-associated drug resistance, we examined the expression of YB-1 and its targeted multidrug resistance genes across multiple conditions. As shown in **Figure S3E**, ATP-binding cassette (ABC) transporters genes (ABCB1, ABCC1, ABCC2), and major vesicle protein (MVP) were upregulated in tumor samples, while their expression was minimal in adjacent tissues. These genes were predominantly expressed at high levels in aneuploid cells but not in diploid cells (**Figure S3F**). Based on predicted cisplatin sensitivity, these genes were more highly expressed in cisplatin-resistant cells rather than cisplatin sensitive genes (**Figure S3G**). Consistently, UMAP density maps showed overlapping high-expression regions of YB-1, ABCB1, ABCC1, ABCC2, and MVP in clusters of malignant cholangiocarcinoma cells (**Figure 1C**, **Figure 3F**). Notably, in gemcitabine-resistant cells, only YB-1 and ABCB1 were co-expressed at high levels (**Figure S3H**). These results indicate that upregulated YB-1 expression may confer broad-spectrum chemotherapy resistance in aneuploid cholangiocarcinoma cells by regulating multidrug resistance transporters.

### YB-1 contributes to cisplatin-induced MDR1 expression

To clarify the role of YB-1 on cholangiocarcinoma chemoresistance, we examined YB-1 expression in the normal immortalized cholangiocyte line MMNK-1, iCCA cell lines (HCCC-9810, HuCC-T1, and CC-SW-1), and eCCA cell lines (TFK-1 and EGI-1). Western blot analysis showed that YB-1 expression in MMNK-1 cells was significantly lower than in all examined cell lines (**Figure 4A**). Notably, the molecular weight of YB-1 was approximately 50 kDa in MMNK-1, HCCC-9810, TFK-1, HUCC-T1, and EGI-1 cells, whereas in CC-SW-1 cells, YB-1 appeared at approximately 40 kDa (**Figure 4A**).

**Figure 4.**
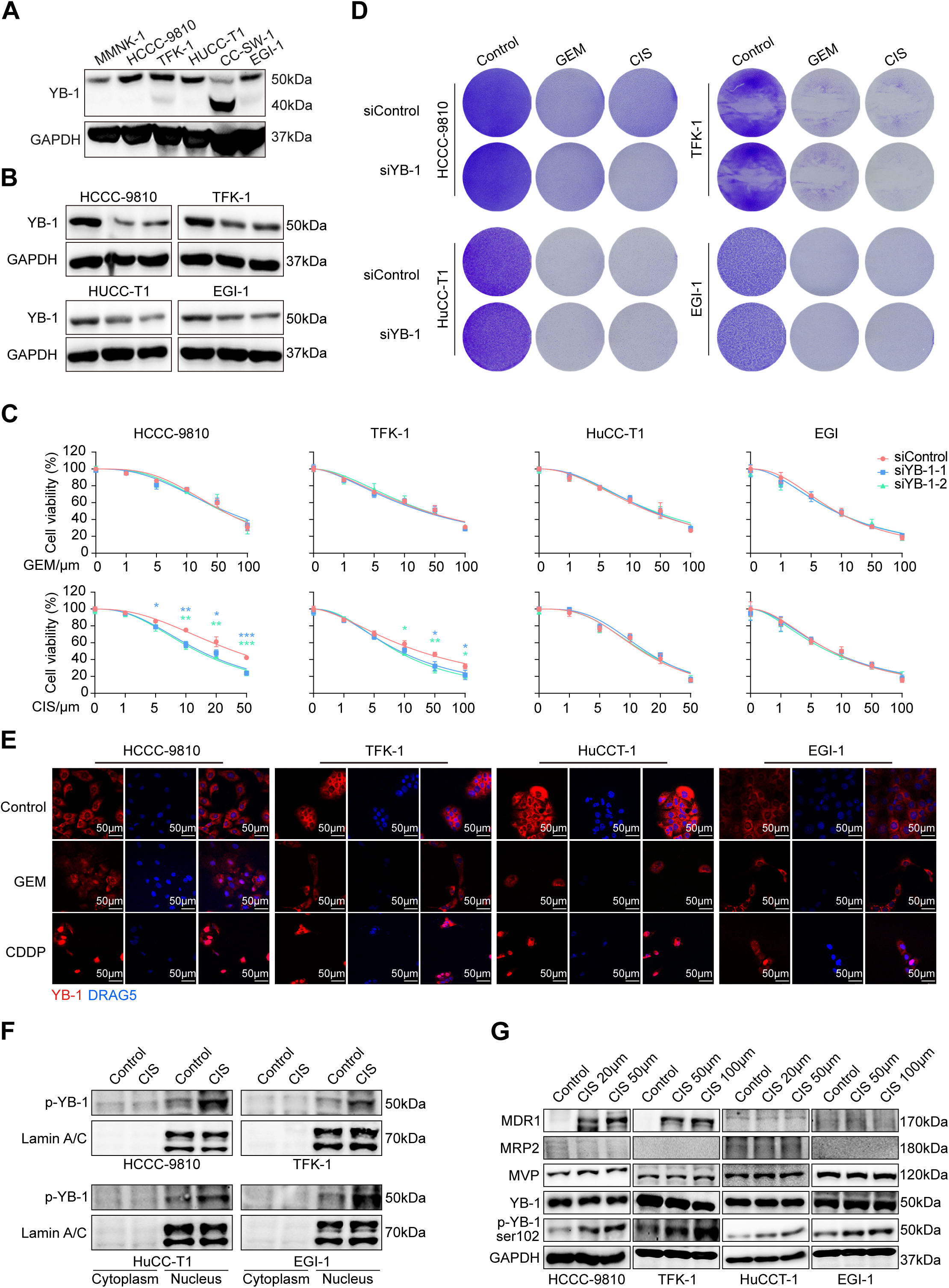
YB-1 expression mediates cisplatin chemoresistance in cholangiocarcinoma cell lines. **(A)** Western blot analysis of YB-1 protein expression across a panel of cholangiocarcinoma cell lines (HCCC-9810, TFK-1, HUCC-T1, CC-SW-1, EGI-1) and the normal cholangiocyte cell line MMNK-1. GAPDH served as a loading control. **(B)** Western blot validation of YB-1 knockdown efficiency in four cholangiocarcinoma cell lines (HCCC-9810, TFK-1, HuCC-T1, and EGI-1). GAPDH serves as the loading control. **(C)** Cell viability assays demonstrating the effect of YB-1 knockdown on cisplatin (CIS) or gemcitabine (GEM) sensitivity. **(D)** Colony formation assays of the indicated cell lines transfected with siControl or siYB-1 and treated with GEM or CIS. Representative crystal violet-stained plates are shown. **(E)** Immunofluorescence showing YB-1 subcellular localization (red) with nuclear counterstaining (DRAQ5, blue) in response to gemcitabine or cisplatin treatment. **(F)** Nuclear-cytoplasmic separation assays analyzing the nuclear translocation of phosphorylated YB-1 cells after cisplatin treatment. Laminin A/C (70 kDa) was used as a nuclear marker and a control for loading nuclear components. **(G)** Western blot analysis of total YB-1, phosphorylated YB-1 (p-YB-1 ser 102), and multidrug resistance-associated proteins (MDR1, MRP2, MVP) following cisplatin treatment with increasing concentrations. GAPDH serves as the loading control.

We next knocked down mRNA and protein expression of YB-1 in two iCCA cell lines (HCCC-9810, HuCC-T1) and two eCCA cell lines (TFK-1, and EGI-1) using two independent siRNA constructs (siYB-1-1 and siYB-1-2) (**Figure S4A, Figure 4B**). Cell viability analyses showed that YB-1 knockdown did not affect cell viability in any of four types of CCA cells treated with gemcitabine (**Figure 4C**). However, silencing YB-1 increased the sensitivity of HCCC-9810 and TFK-1 cells to cisplatin. In these two CCA cells, high concentrations of cisplatin significantly increased cell death following YB-1 knockdown (**Figure 4C**). In contrast, YB-1 silencing did not affect cisplatin response in HuCC-T1 and EGI-1 cells (**Figure 4C**), suggesting a cell context-dependent mechanism of chemoresistance. Consistent with these findings, crystal violet staining showed that YB-1 knockdown enhanced cisplatin-induced cell death in HCCC-9810 and TFK-1 cells, but not in HuCC-T1 and EGI-1 cells (**Figure 4D**). YB-1 knockdown did not affect gemcitabine-induced cell death in any of the examined cell lines (**Figure 4D**).

As a transcription factor, YB-1 translocates to the nucleus upon phosphorylation at serine 102^38^. To examine YB-1 subcellular localization, we performed immunofluorescence staining in HCCC-9810, TFK-1, HuCC-T1 and EGI-1 cells treated with gemcitabine or cisplatin. Without stimulations, YB-1 was primarily localized in the cytoplasm in all examined cell lines (**Figure 4E**). Gemcitabine treatment induced minimal nuclear translocation of YB-1 in HCCC-9810 cell only (**Figure 4E**). In contrast, cisplatin treatment triggered marked nuclear translocation of YB-1 in all examined cell lines (**Figure 4E**). Nucleocytoplasmic fractionation further confirmed that cisplatin significantly increased p-YB-1 levels and promoted its nuclear accumulation (**Figure 4F**). These results suggest that chemotherapy, particularly cisplatin, induces nuclear localization of YB-1 in cholangiocarcinoma cells.

Next, we examined the effects of YB-1 on the expression of multidrug resistance–associated genes (ABCB1, ABCC1, ABCC2, and MVP). qPCR analysis revealed that cisplatin treatment dose-dependently induced ABCB1 mRNA expression in HCCC-9810 and TFK-1 cell lines but not in HuCC-T1 or EGI-1 cells (**Figure S4A**). In contrast, ABCC2 and MVP were induced in all four cell lines following cisplatin treatment, whereas ABCC1 expression decreased in HCCC-9810 and TFK-1 cells and remained unchanged in the other two cell lines (**Figure S4A**).

Western blot analysis further showed that cisplatin treatment upregulated p-YB-1 levels in all examined cell lines and increased MDR1 (decoded by the ABCB1 gene) protein expression in HCCC-9810 and TFK-1 cells only (**Figure 4G**). MRP2 (encoded by the ABCC2 gene) protein was undetectable by Western blotting in these cells, and the expression of YB-1 and MVP was not altered by cisplatin treatment (**Figure 4G**). These findings indicate that cisplatin activates YB-1 and induces MDR1 expression in a subset of CCA cells, thereby contributing to chemoresistance.

### YB-1 promotes transcriptional activation of MDR1

To determine how YB-1 regulates MDR1 expression, we examined MDR1 levels in cholangiocarcinoma cells following YB-1 knockdown. Silencing YB-1 significantly reduced cisplatin-induced MDR1 protein and ABCB1 mRNA expression in HCCC-9810 and TFK-1 cells, not in HuCC-T1 or EGI-1 cells (**Figure 5A-B**). In contrast, YB-1 knockdown did not affect MVP protein or mRNA expression (**Figure 5A**), nor did it alter the mRNA expression of ABCC1 or ABCC2 in any of the examined cell lines (**Figure S5**). Immunofluorescence staining further confirmed that YB-1knockdown decreased cisplatin-induced MDR1 expression in HCCC-9810 and TFK-1 cells, whereas no obvious change was observed in HuCC-T1 cells (**Figure 5C**).

**Figure 5.**
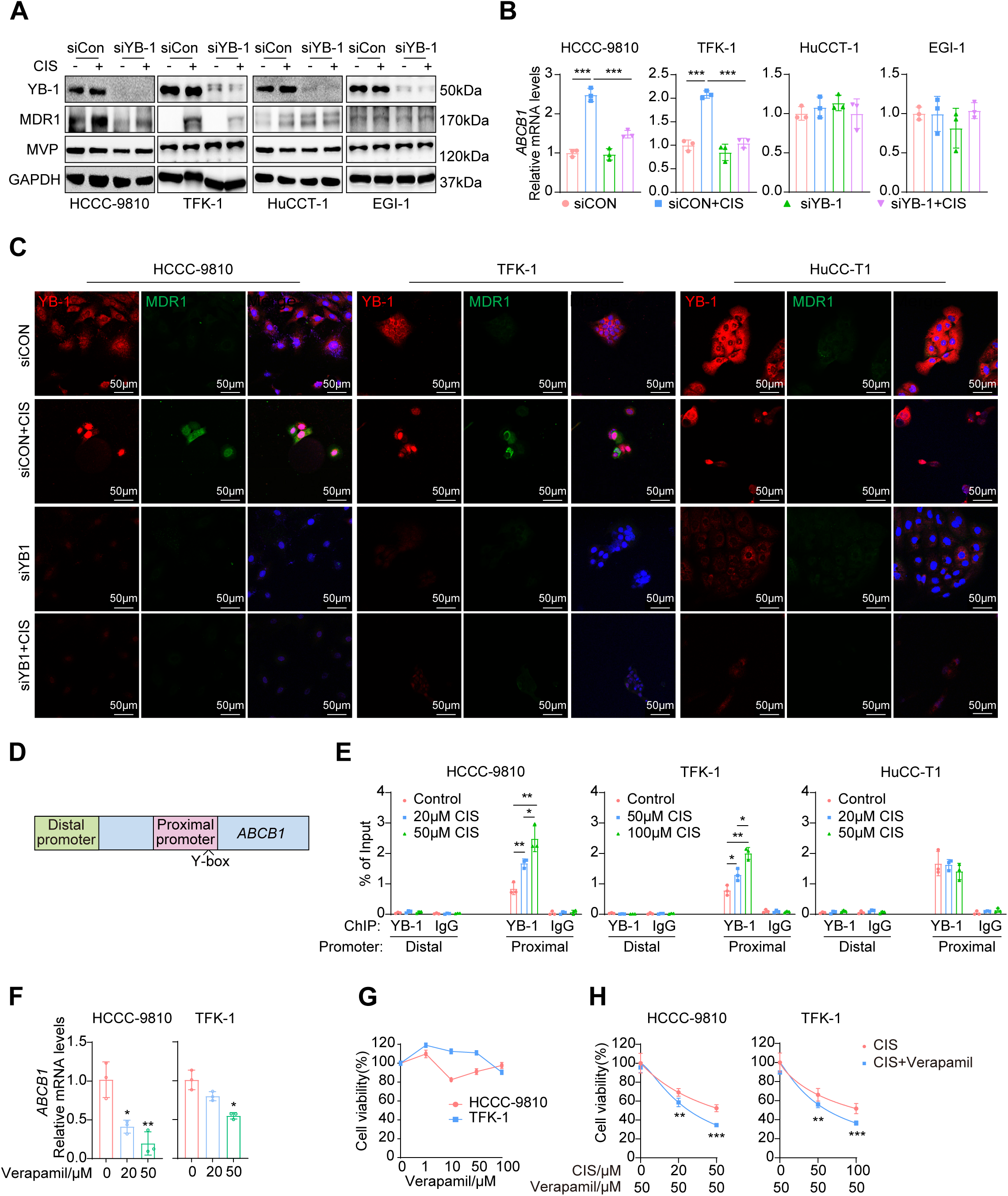
YB-1 transcriptionally upregulates MDR1 expression to promote cisplatin resistance in cholangiocarcinoma. **(A)** Western blot analysis of YB-1, MDR1, MVP, and GAPDH (loading control) in HCCC-9810, TFK-1, HuCCT-1, and EGI-1 cells transfected with siControl (siCon) or siYB-1, with or without cisplatin (CIS) treatment. **(B)** Quantitative RT-PCR analysis of ABCB1 mRNA levels in the indicated cell lines under siCON, siCON+CIS, siYB-1, or siYB-1+CIS conditions. Data are presented as mean ± SEM (n = 3). ***p < 0.001. **(C)** Co-immunofluorescence staining of YB-1 (red) and MDR1 (green) with merged images in HCCC-9810, TFK-1, and HuCC-T1 cells under siCON, siCON+CIS, siYB1, or siYB1+CIS conditions. Scale bar = 50 µm. **(D)** Schematic diagram of the ABCB1 gene promoter structure showing the distal promoter, proximal promoter regions, and Y-box elements. **(E)** Chromatin immunoprecipitation (ChIP) assays in HCCC-9810, TFK-1, and HuCC-T1 cells treated with increasing concentrations of CIS. Enrichment of YB-1 or IgG (negative control) at the distal and proximal ABCB1 promoter regions is shown as percentage of input. *p < 0.05, **p < 0.01. **(F)** Relative ABCB1 mRNA levels in HCCC-9810 and TFK-1 cells treated with the MDR1 inhibitor verapamil (0, 20, or 50 µM), as assessed by quantitative RT-PCR. *p < 0.05, **p < 0.01. **(G)** Cell viability curves of HCCC-9810 and TFK-1 cells treated with increasing concentrations of verapamil. **(H)** Cell viability assays of HCCC-9810 and TFK-1 cells treated with CIS alone or in combination with verapamil (50 µM). **p < 0.01, ***p < 0.001.

To assess whether YB-1 directly regulates ABCB1 transcription, we performed chromatin immunoprecipitation (ChIP) assays in CCA cells with or without cisplatin treatment. Previous studies^39,40^ have shown that the proximal promoter region of the ABCB1 gene contains the YB-1 binding sites (Y-box element) (**Figure 5D**). ChIP-qPCR analysis demonstrated that YB-1 bound to the proximal, but not distal, promoter region in HCCC-9810, TFK-1, and HuCC-T1 cells (**Figure 5E**). Moreover, cisplatin dose-dependently enhanced YB-1 binding to the proximal ABCB1 promoter in HCCC-9810 and TFK-1 cells (**Figure 5E**).

To further clarify the role of MDR1 in cisplatin chemoresistance, we evaluated the effect of the MDR1 inhibitor verapamil on CCA cell viability. Verapamil dose-dependently reduced MDR1 expression in HCCC-9810 and TFK-1 after cisplatin treatment (**Figure 5F**). Although verapamil alone did not affect cell viability (**Figure 5G**), its combination with cisplatin significantly decreased cell viability in these cells (**Figure 5H**), indicating a critical role of MDR1 in cisplatin resistance. Collectively, these results suggested a key role of the YB-1-MDR1 axis in mediating cisplatin resistance in a subset of cholangiocarcinoma cells.

### Cisplatin promotes YB-1 binding to the ABCB1 gene through by inducing promoter demethylation

To further clarify why YB-1-mediated upregulation of MDR1 is only observed in a subset of CCA, we examined the architecture of the ABCB1 proximal promoter. As shown in **Figure 6A**, the ABCB1 proximal promoter contains a CpG island comprising 18 CpG dinucleotides, including a YB-1 binding motif, suggesting a potential link between DNA methylation and YB-1-dependent transcription regulation. Subsequently, we treated HCCC-9810 and HuCC-T1 cells with cisplatin alone or in combination with the DNA methyltransferase inhibitor 5‘-azacytidine (5‘AZA) (**Figure 6B-C**). In both cell lines, 5’AZA alone significantly induced ABCB1 mRNA expression (**Figure 6B-C**). This induction was further enhanced by the combination of cisplatin and 5‘AZA (**Figure 6B-C**). These results suggested that cisplatin may promote ABCB1 transcription by influencing promoter demethylation in CCA cells.

**Figure 6.**
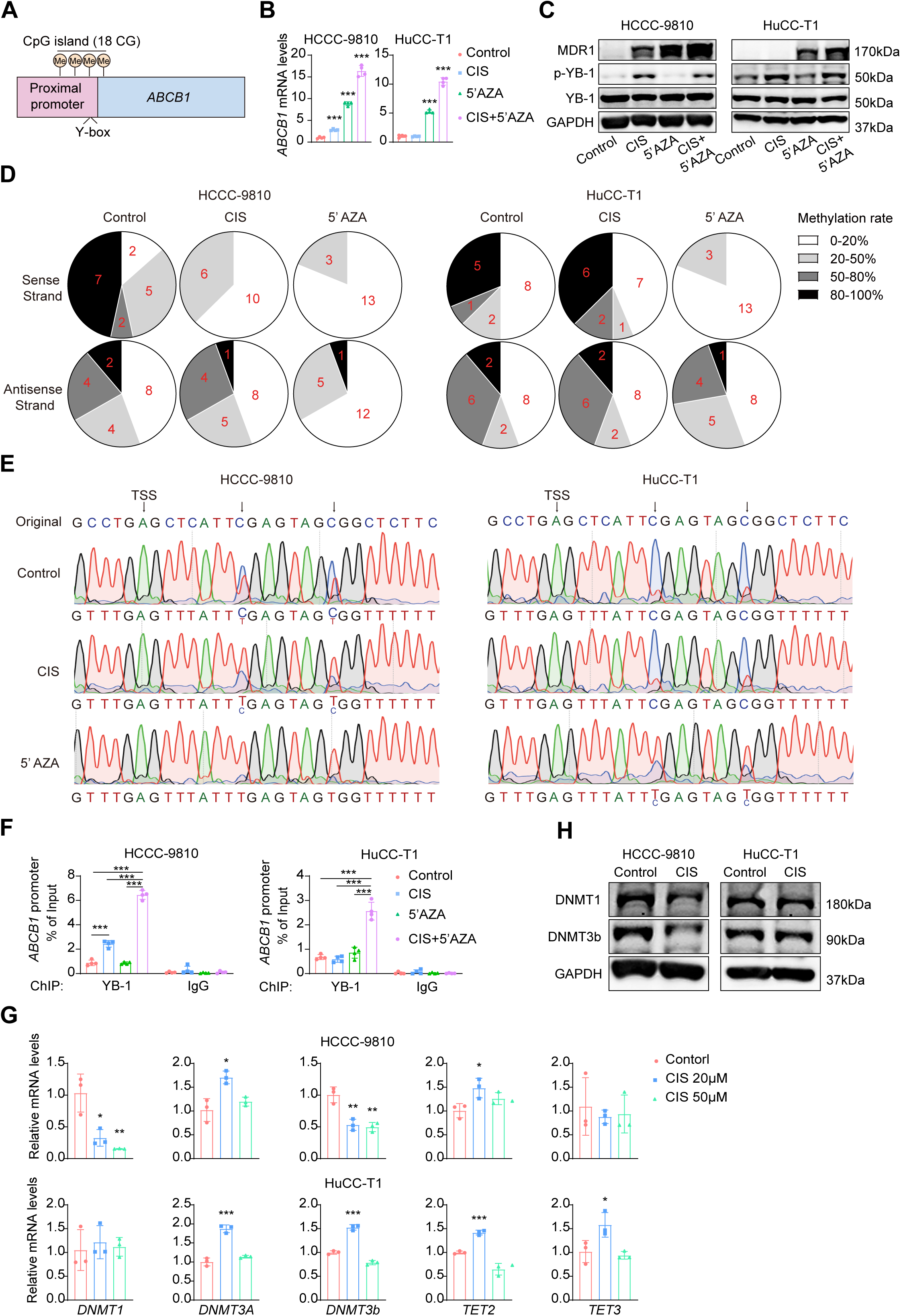
Cisplatin-induced DNA demethylation of the ABCB1 promoter enhances the binding of YB-1. **(A)** Schematic of the ABCB1 proximal promoter region depicting a CpG island containing 18 CG dinucleotides (Me, methylation marks) adjacent to the Y-box binding motif upstream of the transcription start site. **(B)** Quantitative RT-PCR analysis of ABCB1 mRNA levels in HCCC-9810 and HuCC-T1 cells treated with control, CIS, 5’-azacytidine (5’AZA), or the combination CIS+5’AZA. Data are mean ± SEM (n = 3). ***p < 0.001. **(C)** Western blot analysis of MDR1, phospho-YB-1 (p-YB-1), total YB-1, and GAPDH in HCCC-9810 and HuCC-T1 cells under the same four treatment conditions as in (B). **(D)** Bisulfite sequencing analysis comparison of the DNA methylation at the ABCB1 promoter CpG island in HCCC-9810 and HuCC-T1 cells before and after treatment with cisplatin or 5’-azacytidine. The methylation rate is categorized as: white (0-20%), light gray (20-50%), dark gray (50-80%), and black (80-100%). Numbers indicate the count of CpG sites in each methylation category. **(E)** Bisulfite sequencing chromatograms of the ABCB1 proximal promoter region in HCCC-9810 (left) and HuCC-T1 (right) cells under Control, CIS, and 5’AZA treatments. Arrows indicate CpG sites near the transcription start site (TSS); residual cytosine peaks following bisulfite conversion indicate methylated CpGs. **(F)** ChIP assays in HCCC-9810 (left) and HuCC-T1 (right) cells treated with Control, CIS, 5’AZA, or CIS+5’AZA, showing YB-1 enrichment at the ABCB1 promoter as percentage of input. IgG served as a negative control. ***p < 0.001. **(G)** Quantitative RT-PCR analysis of DNMT1, DNMT3A, DNMT3b, TET2, and TET3 mRNA levels in HCCC-9810 (top) and HuCC-T1 (bottom) cells treated with Control, 20 µM CIS, or 50 µM CIS. Data are mean ± SEM (n = 3). *p < 0.05, **p < 0.01, ***p < 0.001. **(H)** Western blot analysis of DNMT1 and DNMT3b protein levels in HCCC-9810 and HuCC-T1 cells treated with Control or CIS. GAPDH serves as the loading control.

We next performed bisulfite sequencing to assess ABCB1 promoter methylation in HCCC-9810 and HuCC-T1 cells. In both cell lines, varying degrees of hypermethylation were observed on both the sense and antisense strands of the ABCB1 promoter CpG island (**Figure 6D**). Treatment with 5’AZA significantly reduced methylation of the sense strand and partially reduced methylation of the antisense strand (**Figure 6D**). Notably, cisplatin treatment reduced methylation of the ABCB1 promoter sense strand only in HCCC-9810 cells (**Figure 6D**). Sequencing peaks analysis further revealed methylation changes at two key CpG sites proximal to the transcription start site (TSS) (**Figure 6E**). In HCCC-9810 cells, methylation levels at both sites were reduced by cisplatin and 5′AZA treatment, whereas in HuCC-T1 cells, demethylation was observed only following 5′AZA treatment (**Figure 6E**).

ChIP–qPCR analysis further showed that 5′AZA treatment alone did not promote YB-1 binding to the ABCB1 proximal promoter in either cell line, whereas cisplatin treatment increased YB-1 binding only in HCCC-9810 cells (**Figure 6F**). However, combined treatment with 5′AZA and cisplatin significantly enhanced YB-1 binding in both cell lines (**Figure 6F**). These results indicate that promoter demethylation facilitates YB-1 binding to the ABCB1 promoter in CCA cells.

Why does cisplatin induce ABCB1 promoter demethylation only in HCCC-9810 cells? To address this question, we assessed the expression of DNA methyltransferases (DNMTs) and Ten-Eleven Translocation (TET) family enzymes. Cisplatin treatment did not significantly alter mRNA expression of DNMT3A, TET2, or TET3 in either HCCC-9810 or HuCC-T1 cells (TET1 expression was undetectable in both cell lines). However, cisplatin dose-dependently reduced both mRNA and protein levels of DNMT1 and DNMT3B in HCCC-9810 cells, but not in HuCC-T1 cells (**Figure 6G-H**), indicating that cisplatin promotes ABCB1 promoter demethylation by suppressing DNMT1 and DNMT3B expression selectively in HCCC-9810 cells.

These results suggest that, in a subset of CCA cells, cisplatin suppresses DNMT1 and DNMT3B expression, thereby inducing demethylation of the ABCB1 promoter CpG island. Demethylation of this CpG island provides a prerequisite for YB-1 binding and subsequent transcriptional activation of ABCB1.

### Cisplatin promotes m⁶A-dependent degradation of DNMT1/3B mRNAs via P-body recruitment

How can cisplatin suppress DNMT1/3B expression in HCCC-9810 cells? The RNA-binding protein HuR is known to stabilize DNMT3b mRNA^41^. A previous study has shown that cisplatin promotes DNMT3B mRNA degradation by disrupting the HuR-DNMT3b mRNA complex in human colorectal carcinoma RKO cells^41^. Therefore, we evaluated the effect of cisplatin on the decay kinetics of DNMT1 and DNMT3B mRNA in HCCC-9810 cells. Cells were treated with the transcriptional inhibitor actinomycin D to block de novo RNA synthesis, and mRNA half-life was subsequently determined. As shown in **Figure 7A**, cisplatin treatment significantly accelerated the decay of DNMT1 and DNMT3B mRNA, indicating reduced transcript stability (**Figure 7A**).

**Figure 7.**
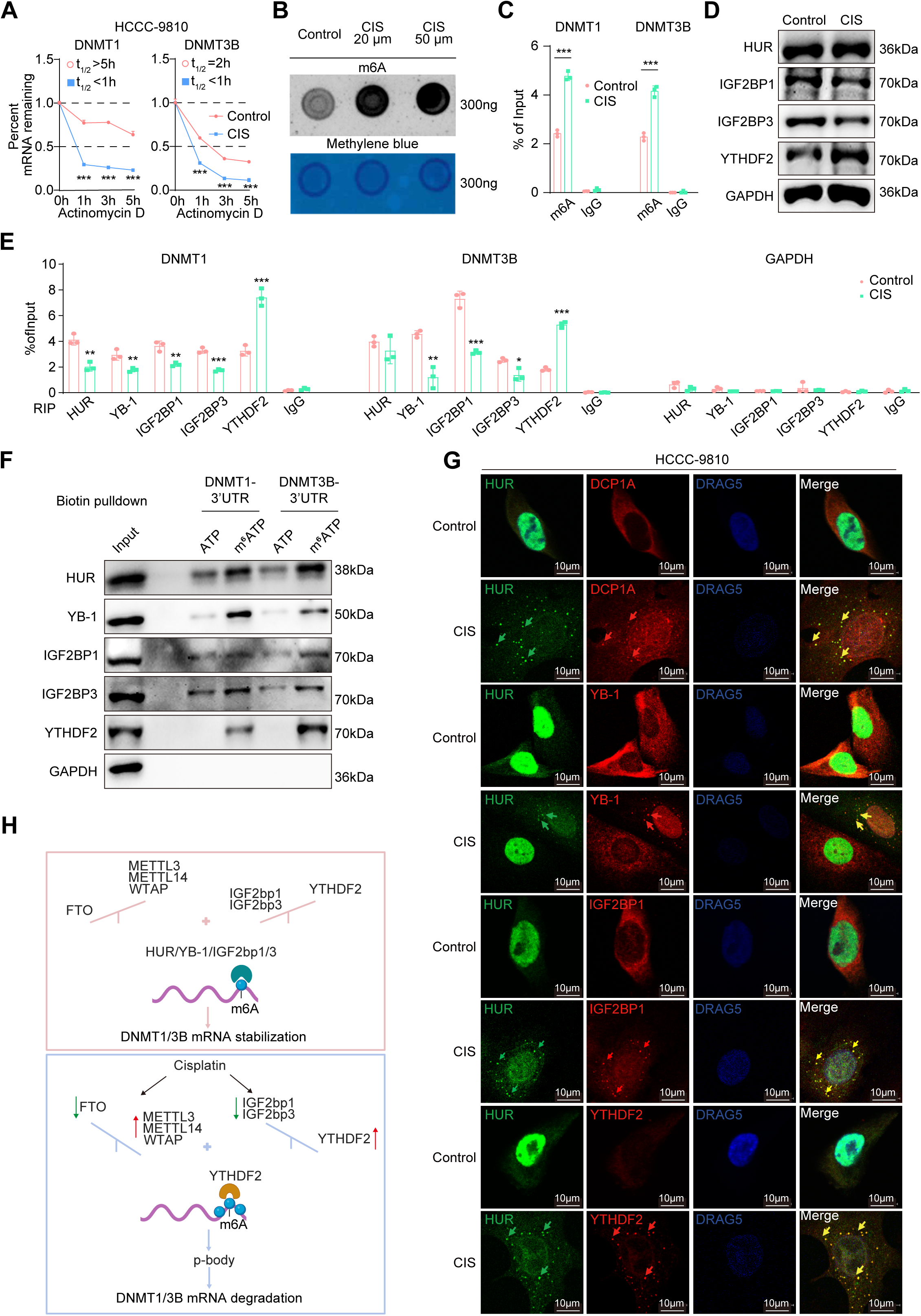
Cisplatin-driven m⁶A remodeling controls DNMT1/3B transcript stability. **(A)** mRNA stability assays for DNMT1 and DNMT3B in HCCC-9810 cells treated with control or CIS, following actinomycin D-mediated transcriptional arrest. Remaining mRNA levels were quantified by RT-PCR at 0, 1, 3, and 5 hours. Estimated half-lives (t½) are indicated. ***p < 0.001. **(B)** Dot blot analysis of global m6A levels in total RNA isolated from HCCC-9810 cells treated with Control, 20 µM CIS, or 50 µM CIS. Methylene blue staining of the same membrane confirmed equal RNA loading (300 ng per spot). **(C)** m6A RNA immunoprecipitation (MeRIP) assays in HCCC-9810 cells (Control vs. CIS) showing enrichment of DNMT1 and DNMT3B transcripts at m6A-modified sites. IgG served as a negative control. Data are mean ± SEM. ***p < 0.001. **(D)** Western blot analysis of RNA-binding proteins HUR, IGF2BP1, IGF2BP3, and YTHDF2 in HCCC-9810 cells treated with Control or CIS. GAPDH served as a loading control. **(E)** RNA immunoprecipitation (RIP) assays assessing binding of HUR, YB-1, IGF2BP1, IGF2BP3, and YTHDF2 to DNMT1, DNMT3B, and GAPDH transcripts in HCCC-9810 cells under Control or CIS conditions. Data are presented as percentage of input ± SEM. *p < 0.05, **p < 0.01, ***p < 0.001. **(F)** Biotin pulldown assays using the 3’UTRs of DNMT1 and DNMT3B transcripts with unmodified (ATP) or m6A-modified (m6ATP) probes. Bound fractions were immunoblotted for HUR, YB-1, IGF2BP1, IGF2BP3, YTHDF2, and GAPDH. **(G)** Co-immunofluorescence imaging in HCCC-9810 cells under Control or CIS conditions. HUR was co-stained with DCP1A (P-body marker), YB-1, IGF2BP1, or YTHDF2, with DRAG5 nuclear counterstain. Yellow arrows indicate co-localization foci. Scale bar = 10 µm. **(H)** Schematic model illustrating the cisplatin-induced m6A reader switch governing DNMT1/3B mRNA stability.

N6-methyladenosine (m^6^A) modification, characterized by the consensus RRACH motif (R=A or G; H=A, C, or U), is the most prevalent internal modification in mRNA, particularly enriched in the 3’UTR (**Figure S6A**). Sequence analysis identified multiple putative m^6^A motifs within the 3‘UTRs of DNMT1 and DNMT3b mRNA (**Figure S6B**). To examine whether m^6^A modification contributes to cisplatin-induced DNMT1/3B mRNA degradation, we examined the expression of m^6^A writers (METTL3, METTL14, and WTAP) and the eraser FTO in cisplatin-treated HCCC-9810 cells. Cisplatin treatment dose-dependently upregulated METTL3, METTL14, and WTAP mRNA expression while downregulating FTO (**Figure S6C**). Consistent with these findings, m^6^A dot blot analysis revealed a dose-dependently increase in global m^6^A levels following cisplatin treatment (**Figure 7B**). Furthemore, m^6^A RNA immunoprecipitation (m^6^A-RIP) assays demonstrated enhanced m^6^A modification of DNMT1/3B mRNAs (**Figure 7C**).

m^6^A modification regulates mRNA stability through reader proteins, including HuR and IGF2BP1/2/3, which promote stabilization, and YTHDF2, which facilitate mRNA decay^41–43^. We therefore examined the expression of these m^6^A readers. Cisplatin treatment did not significantly alter HUR or IGF2BP2 expression but dose-dependently reduced IGF2BP1 and IGF2BP3 levels (**Figure 7D**, **Figure S6D**). In addition, YTHDF2 expression was progressively increased upon cisplatin exposure (**Figure 7D**, **Figure S6D**).

RNA immunoprecipitation assays further showed that HuR, IGF2BP1/3, YTHDF2, and YB-1 bind to DNMT1/3B mRNAs under basal conditions (**Figure 7E**). Cisplatin treatment markedly reduced the binding of HuR, IGF2BP1/3, and YB-1, while significantly enhancing YTHDF2 association (**Figure 7E**). To directly assess the effect of m^6^A on reader binding specificity, in vitro biotin-pulling assays were performed using synthetic RNA probes corresponding to the 3′UTR of DNMT1 and DNMT3B, containing either unmodified adenosine (ATP) or N6-methyladenosine (m^6^ATP). HUR, YB-1, IGF2BP1, and IGF2BP3 preferentially bound to m^6^ATP-modified RNA probes, whereas YTHDF2 bounding exclusively to m^6^A-modified RNA (**Figure 7F**). GAPDH showed no differential binding under any conditions, confirming assay specificity (**Figure 7F**). These results suggest that m^6^A modification facilitates a shift in DNMT1/3B mRNA recognition from a stabilizing complexes to decay-promoting complexes.

Many RNA-binding proteins (RBPs) containing intrinsically disordered regions (IDRs) undergo phase transitions to form biomolecular condensates, which are dynamically regulated by RNA^44–46^. The degradation of m^6^A-modified mRNA has been linked to processing body (P-body) via YTHDF2-mediated phase separation^47^.

We therefore hypothesized that cisplatin may promote P-bodies formation in CCA cells. Indeed, HuR, YB-1, IGF2BP1/3 and YTHDF2 all contain predicted IDRs (**Figure S7**). Immunofluorescence analysis revealed that, in untreated HCCC-9810 cells, HuR was predominantly nuclear, whereas YB-1, IGF2BP1, YTHDF2, and the P-body marker DCP1A were diffusely distributed in the cytoplasm presented as scattered spots (**Figure 7G**), indicating minimal cytoplasmic granule formation. Upon cisplatin treatment, HuR partially relocalized to the cytoplasm, while YB-1 showed partially nuclear relocalization. Simultaneously, prominent DCP1A-positive cytoplasmic foci formed, with strong co-localization of IGF2BP1 and YTHDF2 (**Figure 7G**). This redistribution suggests extensive remodeling of cytoplasmic mRNP complexes. Taken together, these findings support a model in which cisplatin promotes the redistribution of DNMT1/3B-associated RNPs from the IGF2BP-containing stabilizing complexes to the YTHDF2-associated decay complexes within P-body, thereby accelerating mRNA degradation.

## Discussion

Recently, several studies have highlighted the role of YB-1 in CCA progression and chemoresistance^48–51^. Consistent with these studies, we found that YB-1 was enriched in aneuploid malignant cholangiocarcinoma cells. Notably, YB-1 expression, particularly its nuclear localization, is strongly associated with poor prognosis and chemotherapy failure in iCCA. Nuclear YB-1 expression positively correlated with tumor invasion, vascular invasion, postoperative metastasis, and advanced disease stage. More importantly, among the nine patients without nuclear YB-1 expression, eight survived longer than 2 years and two remained alive at the end of follow-up (> 5 years). In contrast, among the five patients with nuclear YB-1 expression, four died within 2 years and one died within 3 years after surgery. Furthermore, among 10 patients receiving postoperative gemcitabine and cisplatin chemotherapy, both YB-1 negative patients (100%) survived until the end of follow-up, whereas, only 2 out of 5 patients (40%) with cytoplasmic YB-1 expression and 1 of 3 patient (33.3%) with nuclear YB-1 expression survived. These results suggest that YB-1, especially its nuclear localization, may play a critical role in chemoresistance.

Cancer cells develop cisplatin resistance through several mechanisms, including reduced intracellular drug accumulation, enhanced DNA damage repair, suppression of apoptosis, and metabolic or mitochondrial reprogramming^52^. Reduced cisplatin accumulation is primarily mediated by ATP-binding cassette (ABC) transporters, particularly ABCB1 encoded by MDR1^52^. Similar to other malignancies, CCA cells can actively export cisplatin through MDR1 overexpression^53^. Previous studies have shown that cisplatin-induced MDR1 expression is associated with activation of mTOR signaling^53^. However, because mTOR signaling broadly regulates protein synthesis, MDR1 induction is unlikely to be explained solely by mTOR activation. In gastric and intestinal cancers, YB-1 has been identified as an important regulator of cisplatin resistance through transcriptional activation of ABCB1^21,22,54,55^. Mechanistically, YB-1 forms a complex with p300 that facilitates RNA polymerase II recruitment to the ABCB1 promoter^54^. Whether YB-1 similarly regulates MDR1 expression in CCA has remained unclear.

In this study, by integrating single-cell transcriptomics, clinical cohort analysis, and mechanistic and functional experiments, we identified a previously unrecognized regulatory axis underlying cisplatin resistance in iCCA. Specifically, cisplatin promoted phosphorylation-dependent nuclear translocation of YB-1, thereby activating ABCB1 transcription. Simultaneously, cisplatin induced m^6^A-dependent destabilization of DNMT1 and DNMT3B mRNAs, leading to demethylation of the ABCB1 promoter, which further facilitated YB-1 binding and transcriptional activation of ABCB1.

To date, few studies have investigated the role of m^6^A modification in cisplatin-induced ABCB1 expression in CCA cells. Our findings suggest that cisplatin initiates a coordinated molecular program leading to ABCB1 activation and chemoresistance through four sequential processes. First, cisplatin induces a cellular stress response that enhances m6A methyltransferase activity while suppressing demethylase activity, resulting in increased m6A deposition on specific transcripts, including DNMT1 and DNMT3B. Second, altered m6A modification shifts RNA-binding protein recruitment, with stabilizing readers such as IGF2BP1/3 being replaced by the destabilizing reader YTHDF2, a process further facilitated by cisplatin-induced changes in reader protein expression. Third, DNMT transcripts are recruited to P-bodies for degradation, leading to reduced DNMT protein levels and passive demethylation of genomic loci, including the ABCB1 promoter. Finally, cisplatin-induced phosphorylation promotes YB-1 nuclear translocation, enabling YB-1 to bind the demethylated ABCB1 promoter and activate transcription of the drug efflux transporter. This regulatory program illustrates how cancer cells coordinate epigenetic and transcriptional stress responses to survive therapeutic pressure. The integration of RNA methylation, DNA methylation, and transcription factor activation creates multiple regulatory checkpoints that may allow tumor cells to adapt dynamically to the intensity and duration of chemotherapy exposure.

Importantly, the cisplatin–YB-1–MDR1 regulatory axis does not appear to operate universally across all CCA cell lines. In HuCC-T1 cells, cisplatin treatment did not alter ABCB1 promoter methylation, and YB-1 disruption failed to influence cisplatin-induced changes in cell viability. These findings suggest that alternative resistance mechanisms may predominate in certain CCA subtypes and warrant further investigation. In addition, unlike its role in cisplatin resistance, YB-1 appeared to play a limited role in gemcitabine resistance. Silencing YB-1 did not significantly alter gemcitabine sensitivity in multiple CCA cell lines. One possible explanation is that gemcitabine treatment does not induce YB-1 nuclear translocation in CCA cells.

In summary, our study identifies a previously unrecognized regulatory circuit linking RNA methylation, DNA methylation, and transcription factor activation in cisplatin resistance in iCCA. Through integration of single-cell transcriptomics, clinical analyses, and mechanistic studies, we demonstrate that YB-1 functions as a central regulator of cisplatin resistance by promoting ABCB1 transcription in cooperation with m6A-dependent destabilization of DNMT1 and DNMT3B. Clinically, the association between nuclear YB-1 expression and poor prognosis, including chemotherapy failure, highlights YB-1 as both a prognostic biomarker and a potential therapeutic target. More broadly, our findings emphasize the importance of multi-layered regulatory mechanisms in therapy resistance and suggest that targeting single signaling pathways alone may be insufficient to overcome adaptive resistance programs. Future studies focusing on biomarker-guided patient stratification and combinatorial therapeutic strategies targeting multiple components of this pathway may improve treatment outcomes for this highly aggressive malignancy.

## Methods and Materials

### Patients

We studied 28 iCCA patients who underwent surgery. Basic clinical data were collected and the stgaes of iCCA were measured according to the eighth edition guideline of American Joint Committee on Cancer. After surgery, 10 patients received adjuvant chemotherapy. All of the patients were followed up at least 6 years. Surgical specimens were also obtained from these iCCA patients. The study protocol fulfilled national laws and regulations and was approved by the local Ethics Committees of University Hospital Tübingen, Germany. These patient samples had been investigated in a previous study^56^.

### Data Sources and Preprocessing

The ICC scRNA-seq dataset was retrieved from GEO (GSE138709) and included five tumor samples and three matched adjacent normal liver samples (33,990 cells in total). All analyses were carried out in R using Seurat 4.0. After quality control, normalization and scaling were performed in Seurat using either the LogNormalize procedure or SCTransform, and highly variable genes were selected for downstream analyses. To reduce inter-sample technical variation, an anchor-based integration procedure was applied (FindIntegrationAnchors and IntegrateData) to generate an integrated expression matrix. PCA was performed, and the retained principal components were used to construct a shared nearest-neighbor graph for graph-based clustering (FindClusters). UMAP^34^ was used to obtain two-dimensional representations of the integrated cellular landscape. Cell-type identities were assigned using marker-gene references from PanglaoDB and CellMarker and supported by published literature.

### Cholangiocyte Sub-clustering and CNV Inference

Cholangiocytes were extracted from the integrated dataset (tumor and adjacent samples) and reanalyzed. PCA and UMAP were performed, followed by unsupervised clustering to define subpopulations within the biliary epithelial compartment. Chromosome-level copy-number variation was inferred from the scRNA-seq expression profiles using inferCNV^35^, with T cells from adjacent tissues used as an internal diploid reference and gene coordinates mapped to hg38. InferCNV was run with default settings, including denoising, and cells were classified as diploid or aneuploid based on the inferred copy-number profiles. The diploid and aneuploid populations were visualized on the UMAP map, and marker-gene annotation based on published studies was used to corroborate the inferCNV-based classification.

### Pseudotime Trajectory Analysis

Trajectory inference and pseudotime analysis were performed using Monocle3 (Trapnell laboratory) on the re-clustered cholangiocyte subset. A principal graph (PCA) was constructed in the reduced-dimensional space, and cells were ordered along pseudotime accordingly. Diploid (non-malignant) cholangiocytes were used as the root population for pseudotime ordering.

### Transcriptional Regulon Analysis (SCENIC and GSVA)

SCENIC^36^ was applied to the single-cell expression matrix to infer gene regulatory networks and quantify transcription factor activity. Candidate target genes were identified from co-expression relationships, and regulons were refined by cis-regulatory motif enrichment analysis. Regulon activity was quantified at the single-cell level using AUCell to obtain AUC-based regulon activity scores, and the activity of the YB-1 regulon was compared between predefined cell groups. In parallel, GSVA was performed using the YB-1 regulon target gene set inferred by SCENIC to compute per-cell enrichment scores, providing an additional readout of YB-1-related transcriptional output.

### Drug Sensitivity Prediction (oncoPredict Analysis)

Drug sensitivity was inferred at single-cell resolution using oncoPredict^37^ with GDSC as the reference. Predicted IC50 values for gemcitabine and cisplatin were calculated from cholangiocyte expression profiles, and cells were classified as predicted sensitive or predicted resistant according to the within-cohort distribution of predicted IC50 values.

### Cell culture

Intrahepatic cholangiocarcinoma (iCCA) cell lines HCCC-9810 and extrahepatic cholangiocarcinoma (eCCA) cell line TFK-1 were cultured in RPMI-1640 (Gibco) medium. iCCA cell lines HuCC-T1 and CCSW-1, eCCA cell line EGI-1, and immortalized human cholangiocyte cell line MMNK1 were cultured in DMEM (Gibco) medium. All the mediums were supplemented with 10% foetal bovine serum (FBS) (Invitrogen) supplemented with 10% FBS (Invitrogen), 4mM L-glutamine (Lonza) and 100 U/mL penicillin/streptomycin (Biochrom KG). Cells were culture in 37 °C and with 5% CO2 and underwent starvation without FBS overnight before treatment with Cisplatin (Sigma), Gemcitabine (Sigma), Verapamil (TOCRIS), 5’Azacytidin (Sigma) and/or Actinomycin D (Thermo scientific).

### Immunohistochemistry (IHC)

Immunohistochemical staining was performed as previously described^57^. Briefly, tissue sections were dewaxed through a graded series of ethanol and washed with phosphate-buffered saline (PBS). Sections were then immersed in 10 mM sodium citrate buffer (pH 6.0), and antigen retrieval was performed using a microwave oven. After cooling to room temperature, sections were incubated with a peroxidase inhibitor (Dako) for 1 hour. The primary antibody was applied, and sections were incubated overnight at 4°C. The following day, after washing with PBS, sections were incubated with EnVision peroxidase (Dako) for 1 hour at room temperature. Finally, the signal was developed with diaminobenzidine (DAB) for 5 minutes.

### Immunofluorescence

Immunofluorescence was performed as described previously^57^. Cells were fixed with 4% PFA for 10 mins, washed by PBS for three times, permeabilized with 0.25% Triton X-100 in PBS for 10 mins, washed by PBS for three times and were incubated by PBST with 22.52 mg/mL glycine and 1% BSA for 1 hour at room temperature. Subsequently, the cells were incubated with 10 μ g/mL YB-1, MDR1, IGF2bp1, YTHDF2, DCP1a and/or HuR antibodies in 1% BSA at 4°C overnight. Next day, cells were washed by PBS-Tween for three times, incubated with 2µg/mL Alexa Fluor® 488 and/or Alexa Fluor® 555 secondary antibodies (Abcam) for 1 hour at room temperature, washed by PBS-Tween for three times and incubated with 5μM DRAQ5 (Cell signaling) for 5 mins. Images were scanned under a Leica confocal microscope (TCS SPE).

### RNA extraction and quantitative real time-PCR

Total RNA was extracted from cells with TRIzol reagent according to the manufacturer’s instructions. cDNA was synthesized using RevertAid H Minus Reverse Transcriptase (Thermo Fischer Scientific). The qRT-PCR assay was performed using POWRUP SYBR MASTER MIX (Life Technologies) by a StepOnePlus Real-time PCR instrument (Applied Biosystems). All the qPCR primers were in the Table S1.

### Gene knockdown

Non-targeting siRNA (CCUACGCCACCAAUUUCGU) and siRNA targeting YB-1(1: GCAGACCGUAACCAUUAUA and 2: GUAAGGAACGGAUAUGGUU) were transfected into cells with Lipofectamine RNAiMAX (Invitrogen) according to the manufacturer’s instruction. After 24h transfection, Cells received different treatments.

### Western blotting

Immunoblot experiments were performed as previously described^57^. Cell protein lysates were treated with RIPA buffer containing protease and phosphorylase inhibitors. Equal amounts of protein (40 µg) were separated by 8% SDS-PAGE electrophoresis and transferred to nitrocellulose membranes. The membranes were blocked with 5% bovine serum albumin (BSA) at room temperature for 1 hour, and then incubated overnight with primary antibodies at 4°C. After washing three times with PBS-Tween, the membranes were incubated with the corresponding secondary antibody. After washing three more times, the membranes were developed with chemiluminescent substrate.

### Nuclear/Cytoplasmic Protein Extraction

Total cell protein was extracted by RIPA buffer with protease and phosphatase inhibitors. Nuclear and Cytoplasmic protein were extracted using the NE-PER™ Nuclear and Cytoplasmic Extraction kit (Thermo Fischer Scientific). 40µg protein was proceeded to western blotting.

### Cell viability assay

After treatment with chemical compounds 48h, cells were incubated with 5 mg/mL 3-(4,5-dimethylthiazol-2-yl)-2,5-diphenyl tetrazolium bromide (Sigma-Aldrich) for 5 h. Following removal of the supernatant, cells were incubated with 100 µL dimethyl sulfoxide for 20min. Absorbance was measured at 570 nm.

### Crystal violet staining

Cells were seeded at a density of 5 × 10⁴ cells per well in 6-well plates and cultured to the desired confluence, then treated with cisplatin or gemcitabine for 48 hours. Following treatment, the culture medium was removed, and the wells were gently washed twice with PBS. Cells were then fixed with ice-cold 100% methanol for 10 minutes and subsequently stained with 0.1% crystal violet at room temperature for 15 minutes. Excess dye was removed by washing the plates several times with distilled water, and the plates were air-dried before imaging.

### Bisfulte sequencing

The genomic DNA was extracted with a QIAamp DNA Micro Kit (Qiagen) and bisulfite converted using an EpiTect Bisulfite Kit (Qiagen). Converted gDNA was performed PCR with KAPA2G Robust HotStart PCR-Kit (Sigma). The PCR-amplified products were subjected to Sanger sequencing. The primers were in the Table S1.

### RNA stability assay

Cells were seeded in 12-well plates and allowed to adhere overnight. The following day, cells were treated with cisplatin for 48 hours. Subsequently, actinomycin D was added to the cells, and total RNA was extracted at 0, 1, 3, and 5 hours after actinomycin D treatment. The levels of the target gene were then quantified by qPCR.

### Ribonucleoprotein Immunoprecipitation

50 μl Protein A/G-agarose beads was washed three times with NT2 buffer (50 mM Tris-HCl (pH 7.4), 150 mM NaCl, 1 mM MgCl2, 0.05% NP-40, with 1× protease inhibitor cocktail), incubated with 5μg relevant antibodies (anti-mouse HUR and anti-mouse IgG as control) (Santa Cruz) overnight at 4°C with gentle rotation and then pulled down with polysome lysis (150 mM KCl, 25 mM Tris-HCl (pH 7.4), 5 mM EDTA, 0.5% NP-40, 0.5 mM DTT, with 100 U/ml RNase inhibitor and 1× protease inhibitor cocktail) extracted protein (500μg) overnight at 4°C with gentle rotation. After washing six times with cold NT2 buffer to remove unbound proteins, the beads were subjected to RNA extraction, cDNA synthesis and qPCR.

### Biotinylated RNA pull-down

Reverse-transcribed total RNA was used as the template for PCR amplification (3′UTR of DNMT1 and DMT3B mRNA). All products contained the T7 RNA polymerase promoter sequence in 5′oligonucleotides. The purified PCR-amplified products were used as templates for the synthesis of biotinylated RNA using T7 RNA polymerase and biotin-UTP with ATP or ATP/m6 ATP mix (1:1). The biotinylated transcripts were incubated with streptavidin-coupled dynabeads (Invitrogen) at 4 °C for 2 h and then the mixture was incubated with cell lysates at 4 °C for another 4 h. The beads were washed with TENT (10 mM Tris-HCl (pH 7.5), 1 mM EDTA (pH 8), 0.25M NaCl, 0.1 % Tween-20) buffer three times and eluted with 2× Laemmli SDS sample buffer. The elution samples proceeded to western blotting.

### m^6^A dot blot assay

A total of 300 ng of RNA was denatured at 65°C for 5 minutes and then spotted onto a nitrocellulose membrane, followed by baking at 80°C for 1–2 hours to immobilize the RNA. Total RNA was visualized by staining with 0.02% methylene blue. The membrane was subsequently blocked with 5% non-fat milk in PBST for 1 hour at room temperature and incubated overnight at 4°C with an anti-m⁶A antibody. After three washes with PBST, the membrane was incubated with an HRP-conjugated secondary antibody for 1 hour at room temperature, and signals were detected using enhanced chemiluminescence.

### ChIP-qPCR

Immunofluorescence was performed as described previously^57^. Briefly, Cells were incubated with 1% formaldehyde for 10 minutes for cross-linking and then 125 mM glycine to terminate the reaction for 5 minutes at room temperature. cells were resuspended in lysis buffer and sonicated to obtain around 500 bp of DNA. Immunoprecipitation aliquots were incubated with 5 μg of YB-1 antibody or rabbit IgG overnight at 4 °C. 50 μl Protein A/G agarose beads (Santa Cruz) and 4 μg herring sperm DNA was added to each sample and incubated at 4° C for 4 hours. Then the beads were washed for 5 minutes at 4 °C with low salt wash buffer, high salt wash buffer, LiCl wash buffer, and TE buffer. Subsequently, the beads were suspended in a 120 μl elution buffer. Following the elution samples and the input samples incubating with 5M NaCl and RNase A (Sigma) at 65 °C overnight to reverse the crosslinks, the samples were incubated with proteinase K (Sigma) at 60 °C for 1 hour. All samples were purified using a MinElute PCR Purification Kit (Qiagen) and analyzed by qPCR.

### Statistical analysis

Associations between YB-1 expression and clinicopathological indexes were tested by Chi-square test. For survival analysis, overall survival was censored from the date of surgery to the date of last follow-up; events were defined as death from the tumor. The survival curves were plotted by the Kaplan–Meier method, and compared by the log-rank test. Measurement data were expressed as mean ± standard deviation (SD). Different groups were compared with unpaired Student’s t test or Mann-Whitney U test using the SPSS 12.0 program. P values less than 0.05 were considered significant.

All data were expressed as mean ± standard deviation (SD). Statistical analyses for experimental data were performed using Student’s t-test for comparisons between two groups and one-way ANOVA for multiple group comparisons. Statistical significance was defined as P < 0.05, with significance levels indicated as follows: *, P < 0.05; **, P < 0.01; ***, P < 0.001 and ****, P < 0.0001.

## Author’s contributions

T.L. H.L.W. conceived and designed the project. J.L. collected the patient samples. H.L. and H.L.W. performed pathological evaluation. T.L., C.T., and H.L. undertook experiments. L.T., C.H., and C.T. performed bioinformatic analyses. T.L. and H.L.W. drafted the article. T.L., C.T., L.L., R.L., J.L., J.A.L., M.P.A.E., P.R.C.M., S.D., and H.L.W. discussed the data and edited the article critically.

## Acknowledgments

We thank the Human Tissue and Cell Research Foundation, a nonprofit foundation regulated by German civil law, which facilitates research with human tissue through the provision of an ethical and legal framework for prospective sample collection. We acknowledge the support of the LIMA Live Cell Imaging at Microscopy Core Facility Platform Mannheim (CFPM). We gratefully acknowledge the data storage service SDS@hd supported by the Ministry of Science, Research and the Arts Baden-Württemberg (MWK) and the German Research Foundation (DFG) through grant INST 35/1503-1 FUGG.

## Figure legends

**Figure S1.**
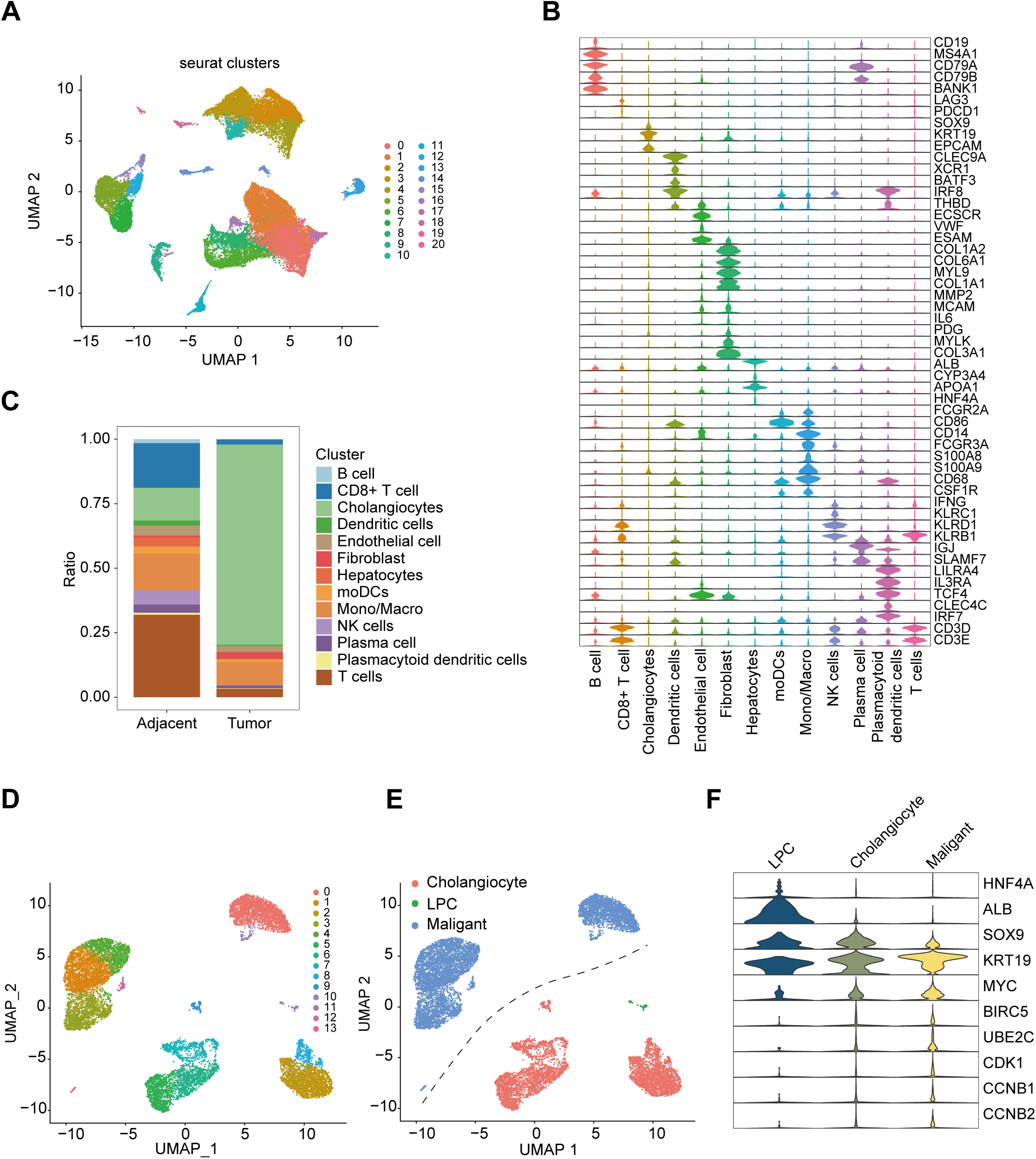
Cellular and epithelial subtype mapping at single-cell resolution. **(A)** UMAP visualization of single-cell RNA sequencing showing 21 distinct cell clusters identified. **(B)** Violin plots displaying the expression patterns of canonical marker genes across different cell types in Figure 1A. **(C)** Stacked bar chart quantifying the relative proportions of each cell type in adjacent versus tumor samples. **(D)** Higher-resolution re-clustering of cholangiocytes further delineated multiple subclusters. **(E)** UMAP visualization of epithelial cells classified into three major subtypes: malignant cells (blue), cholangiocytes (red), and liver progenitor cells (LPC, green). **(F)** Violin plots showing the expression distribution of key marker genes across the three cell types.

**Figure S2.**
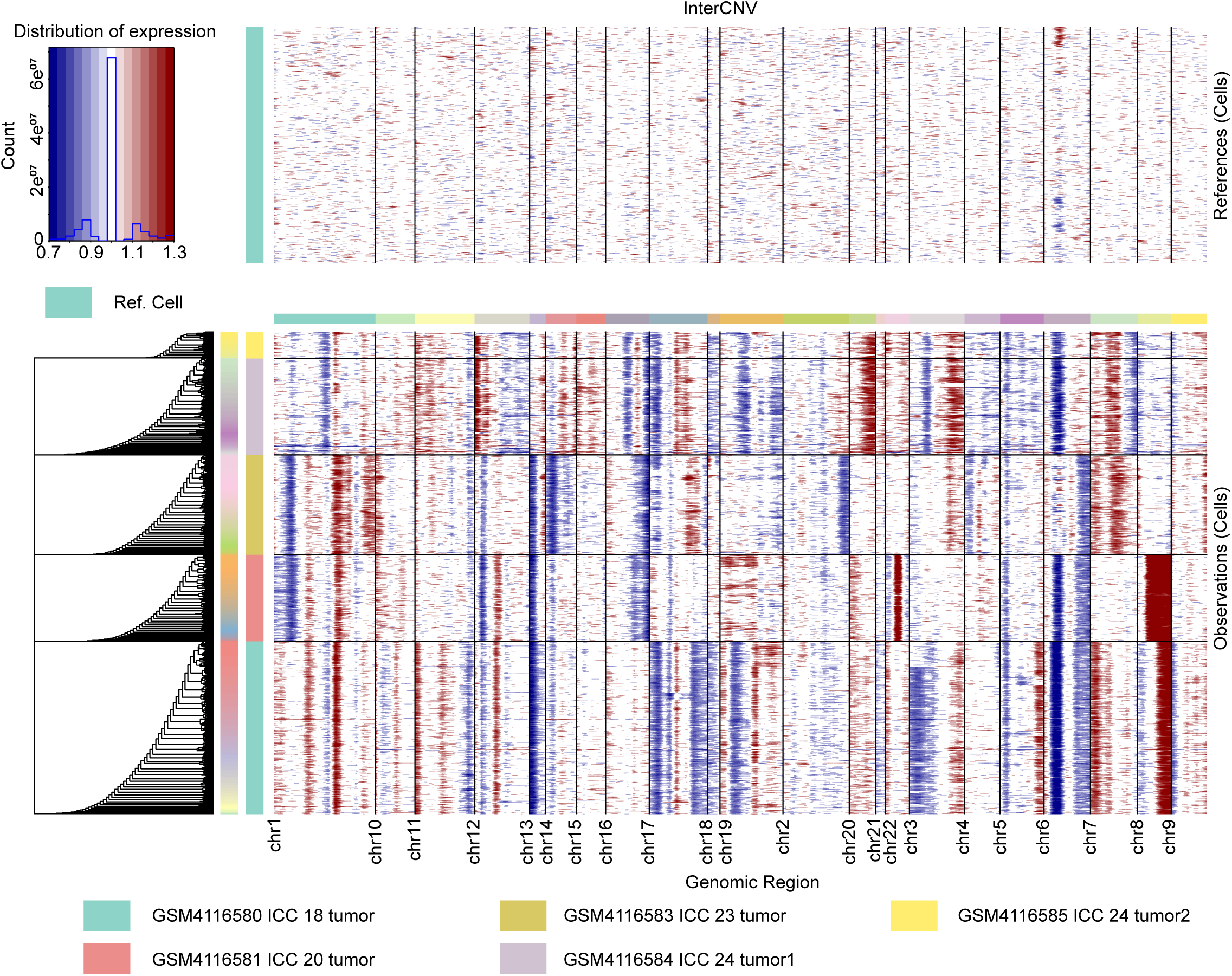
Single-cell CNV landscapes across tumor genomes. Comprehensive copy number variation (CNV) analysis using InferCNV across the genome. Top panel: Heatmap depicting CNV profiles of reference cells, illustrating the distribution of expression levels. Bottom panel: Large-scale CNV heatmap showing chromosomal alterations in individual tumor cells across multiple samples. The color bar indicates CNV status (blue = loss, white = neutral, red = gain). Rows represent individual cells, and columns correspond to genomic regions.

**Figure S3.**
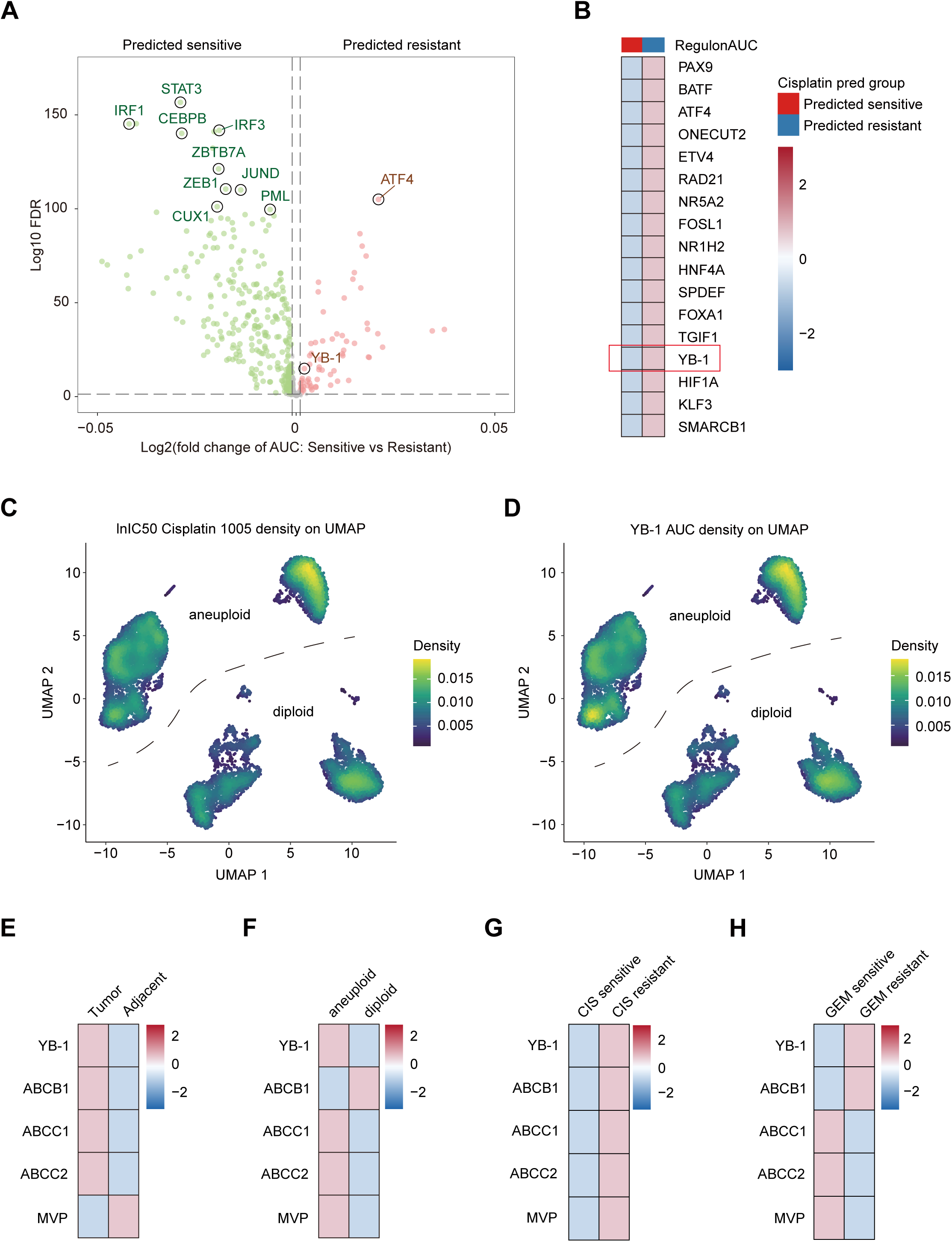
YB-1 regulon activity and chemotherapy resistance across sensitive and resistant cell populations. **(A)** Volcano plot showing differential regulon activity between predicted cisplatin-sensitive (left) and cisplatin-resistant (right) cells. The x-axis represents log2 fold change in regulon AUC scores, and the y-axis shows -log10 of the False Discovery Rate (FDR). **(B)** Heatmap displaying the scaled regulon AUC activity scores for key transcription factors across predicted cisplatin-sensitive (red) and predicted cisplatin-resistant (blue) cell populations. The color scale (blue to red) represents regulon activity levels. **(C-D)** UMAP density plot showing the spatial distribution of predicted cisplatin IC50 values (IntC50_Cisplatin_1005) and YB-1 regulon AUC activity across aneuploid and diploid cell populations. The density values (yellow-green) indicate the degree of cell enrichment with specific IC50 predictions. **(E-H)** Heatmaps illustrating the coordinated expression of YB-1 and chemotherapy resistance–related transporter genes (ABCB1, ABCC1, ABCC2, MVP) across multiple biological contexts, including tumor versus adjacent tissue (**E**), aneuploid versus diploid cells (**F**), cisplatin-sensitive versus cisplatin-resistant cells (**G**), and gemcitabine-sensitive versus gemcitabine-resistant cells (**H**). Normalized expression levels are indicated by the color scale.

**Figure S4.**
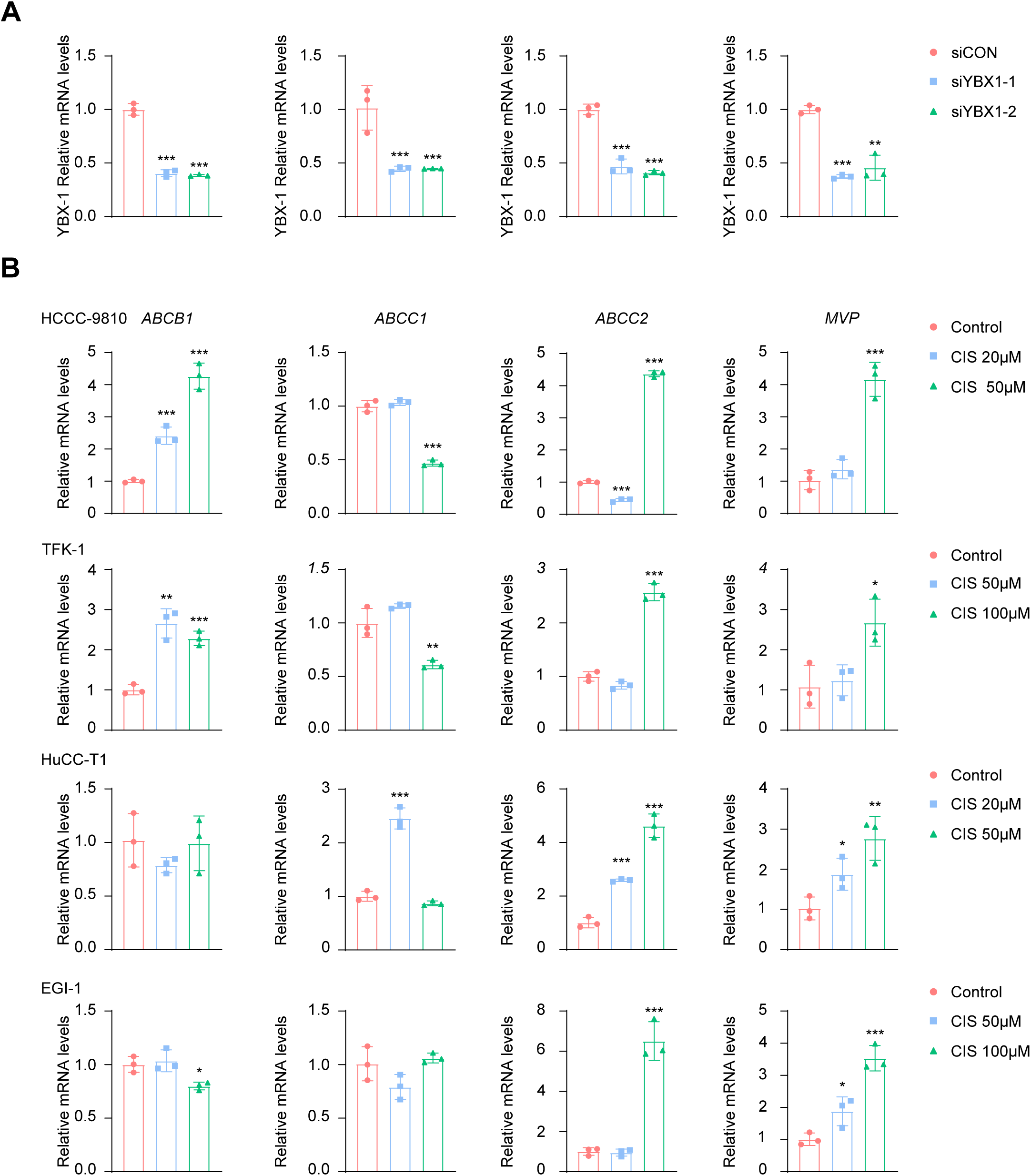
YB-1 knockdown efficiency and cisplatin-Induced differential regulation of multidrug resistance gene expression. **(A)** Quantitative RT-PCR validation of YB-1 (YBX1) mRNA knockdown efficiency in HCCC-9810, TFK-1, HuCC-T1, and EGI-1 cells transfected with siControl (siCON), siYBX1-1, or siYBX1-2. Data are mean ± SEM (n = 3). **p < 0.01, ***p < 0.001. **(B)** Quantitative RT-PCR analysis of multidrug resistance-associated transporter mRNA levels-ABCB1, ABCC1, ABCC2, and MVP-in HCCC-9810, TFK-1, HuCC-T1, and EGI-1 cells treated with increasing concentrations of cisplatin (CIS). Data are mean ± SEM (n = 3). *p < 0.05, **p < 0.01, ***p < 0.001.

**Figure S5.**
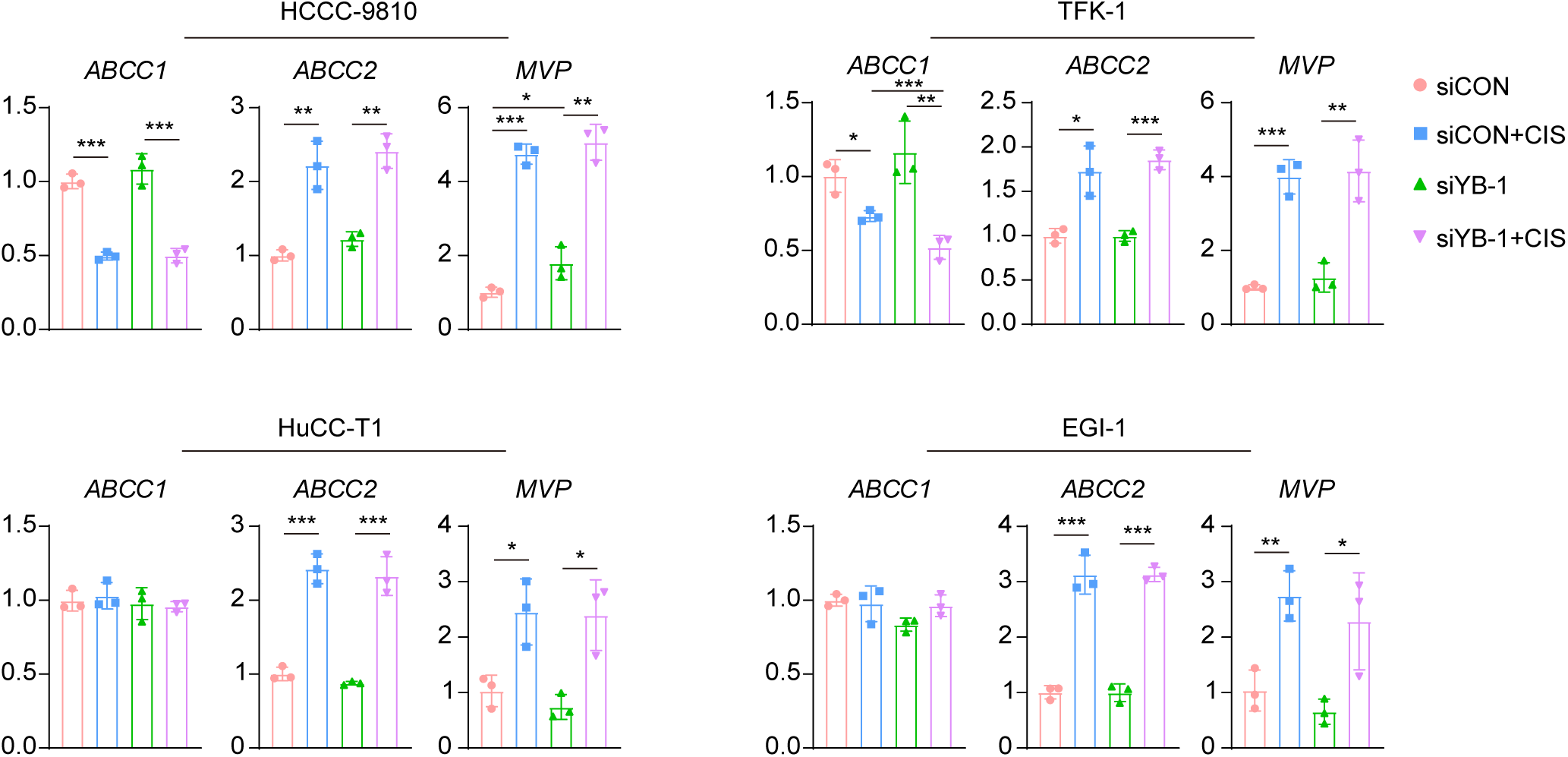
Effect of YB-1 knockdown on the expression of multidrug resistance-associated genes under cisplatin treatment. Quantitative RT-PCR analysis of ABCC1, ABCC2, and MVP mRNA levels in HCCC-9810, TFK-1, HuCC-T1, and EGI-1 cells transfected with siCON or siYB-1, with or without cisplatin (CIS) treatment. Data are mean ± SEM (n = 3). *p < 0.05, **p < 0.01, ***p < 0.001.

**Figure S6.**
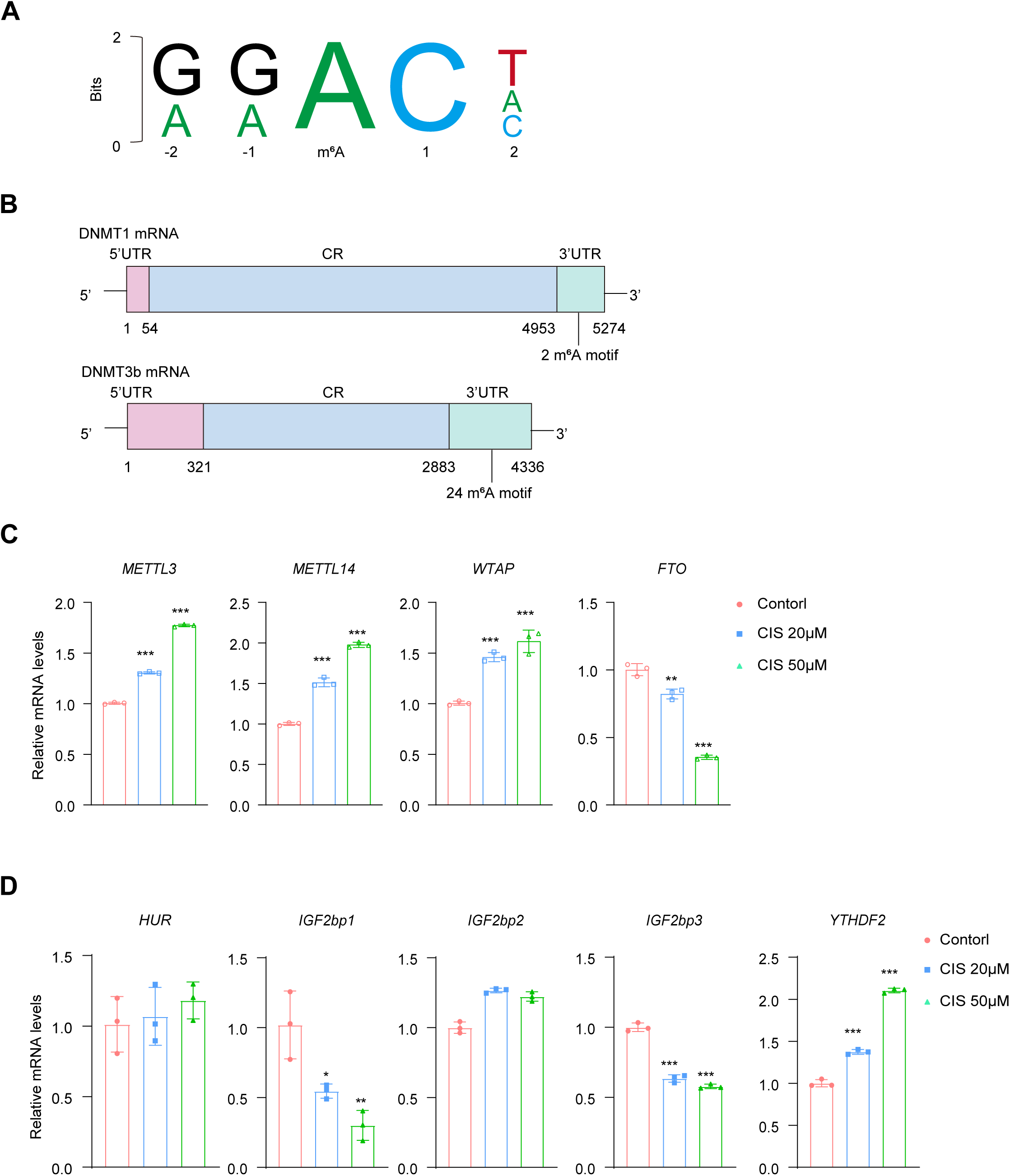
Cisplatin-induced changes in m⁶A writers, erasers and readers mRNAs. **(A)** Sequence marker diagram of the m⁶A (N⁶-methyladenosine) motif. Typical m⁶A modification sites follow the concordant sequence R(G/A)R(G/A)ACH(U/C/A), where the central adenosine (A) (labeled m⁶A) is the methylation site. **(B)** Schematic diagram of the DNMT1 and DNMT3b mRNA structures, showing the distribution of the m⁶A motif. **(C)** Quantitative RT-PCR analysis of m6A writer (METTL3, METTL14, WTAP) and eraser (FTO) mRNA levels in HCCC-9810 cells treated with Control, CIS 20 µM, or CIS 50 µM. Cisplatin dose-dependently upregulated the three m6A writers while significantly downregulating the m6A eraser FTO, indicating a shift toward increased m6A deposition. Data are mean ± SEM (n = 3). **p < 0.01, ***p < 0.001. **(D)** Quantitative RT-PCR analysis of m6A reader mRNA levels-HUR, IGF2BP1, IGF2BP2, IGF2BP3, and YTHDF2-in HCCC-9810 cells treated with Control, CIS 20 µM, or CIS 50 µM. Data are mean ± SEM (n = 3). *p < 0.05, **p < 0.01, ***p < 0.001.

**Figure S7.**
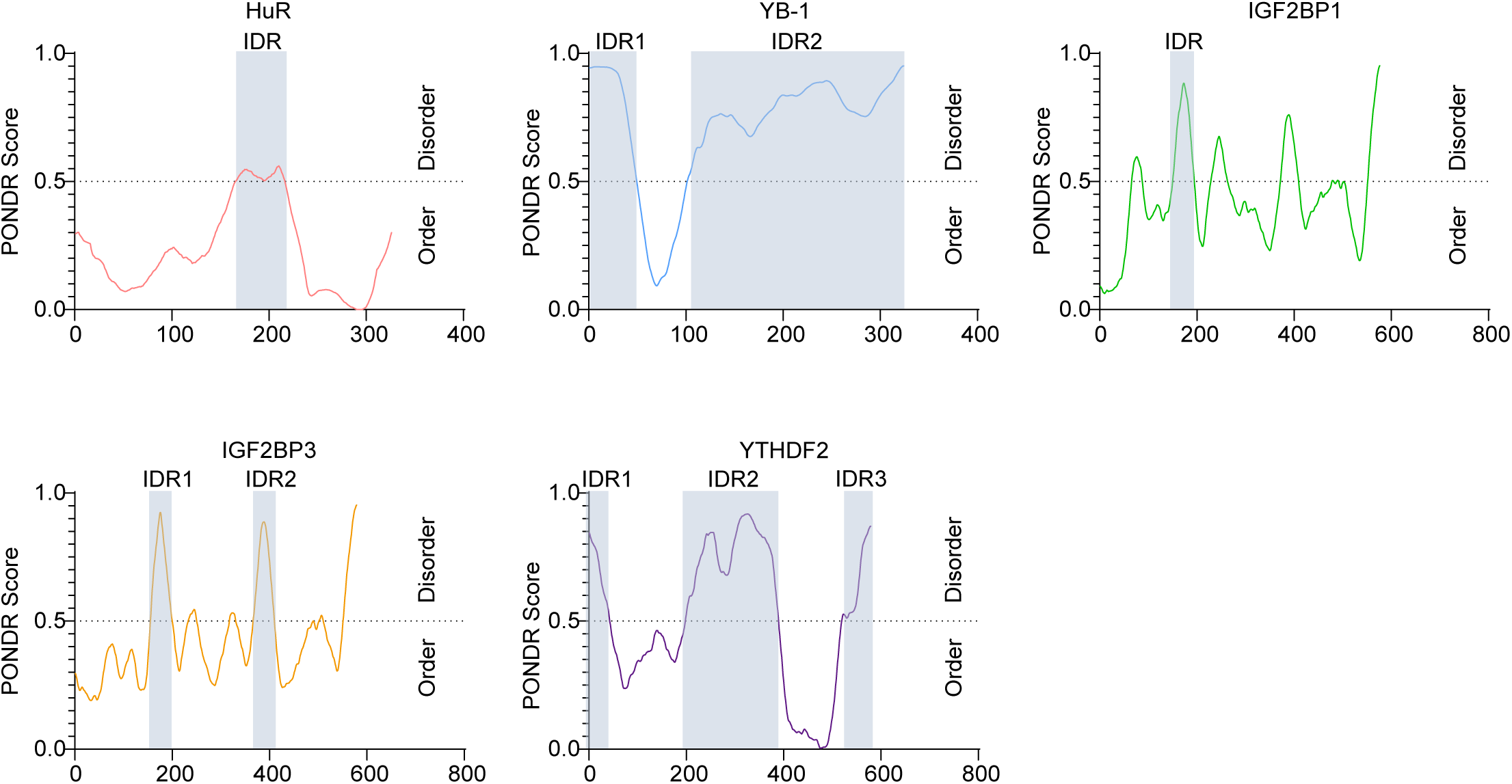
Intrinsically disordered region (IDR) prediction in RNA-binding proteins. PONDR score analysis of RNA-binding proteins HuR, YB-1, IGF2BP1, IGF2BP3 and YTHDF2 sequence reveals the predicted IDRs (shaded region). PONDR (Predictor of Natural Disordered Regions) scores above 0.5 (dashed line) indicate predicted disorder. Shaded regions denote IDRs that exceed the disorder threshold. X-axis represents amino acid position; Y-axis represents disorder prediction score (0-1).

## Key resource table

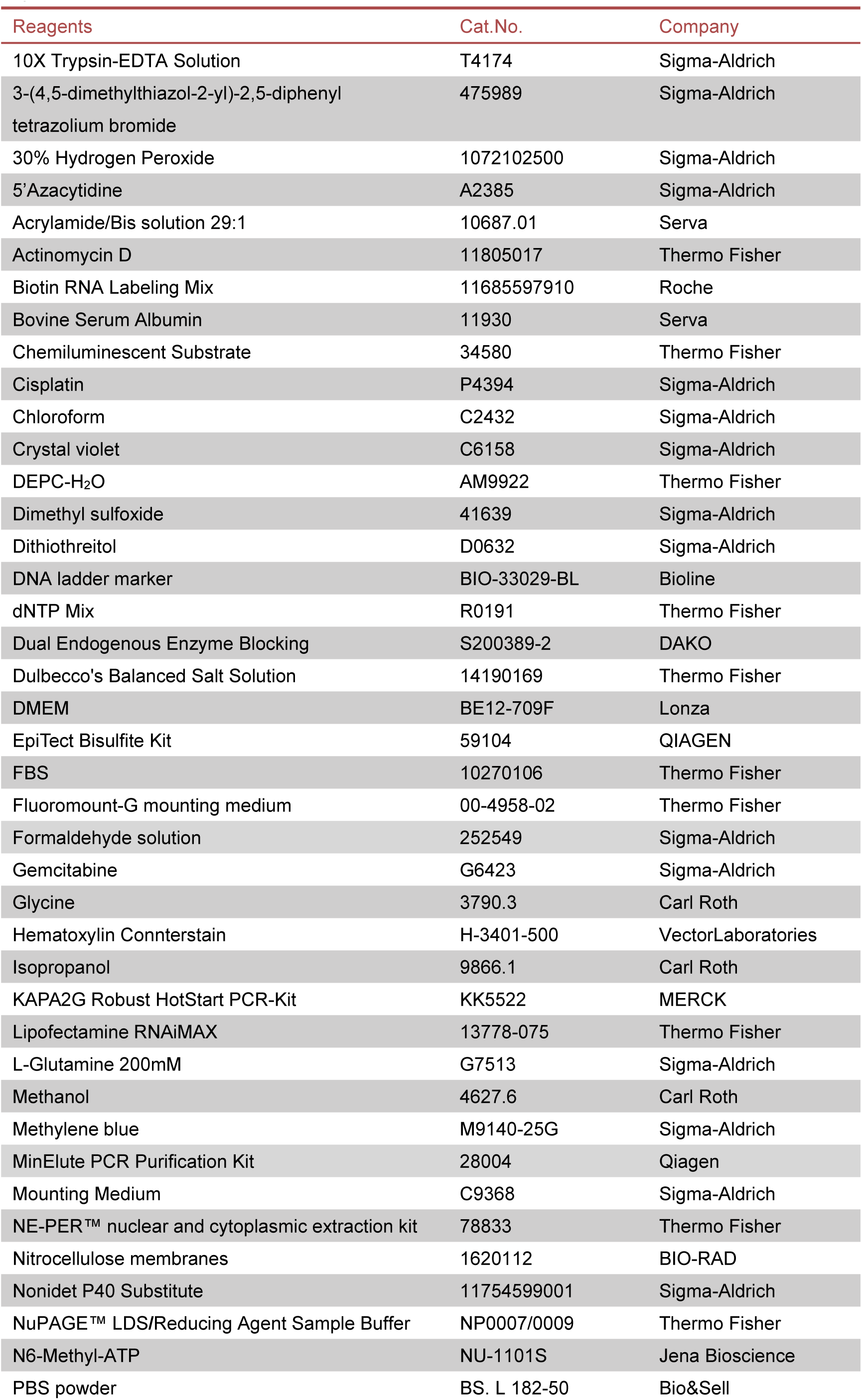
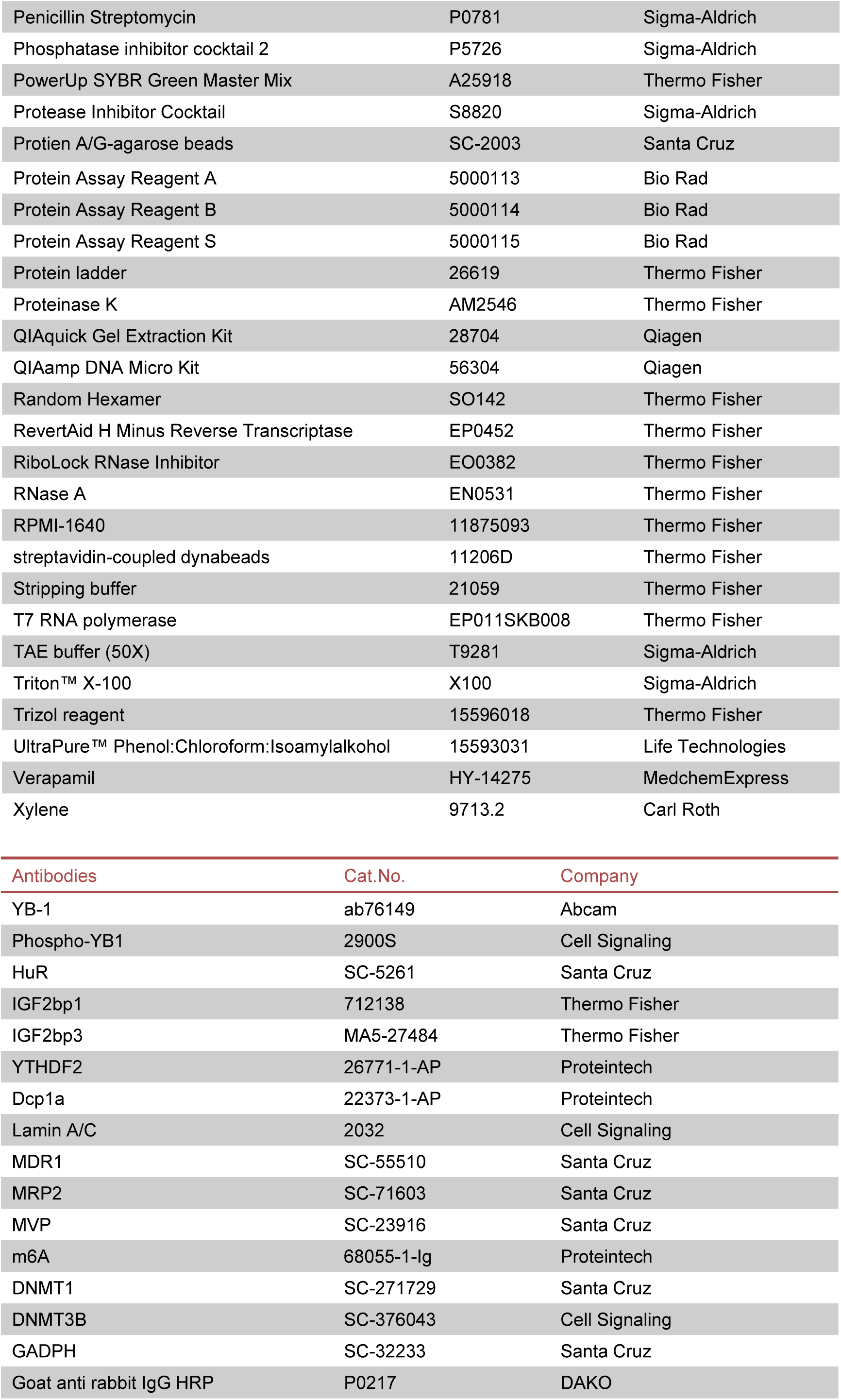
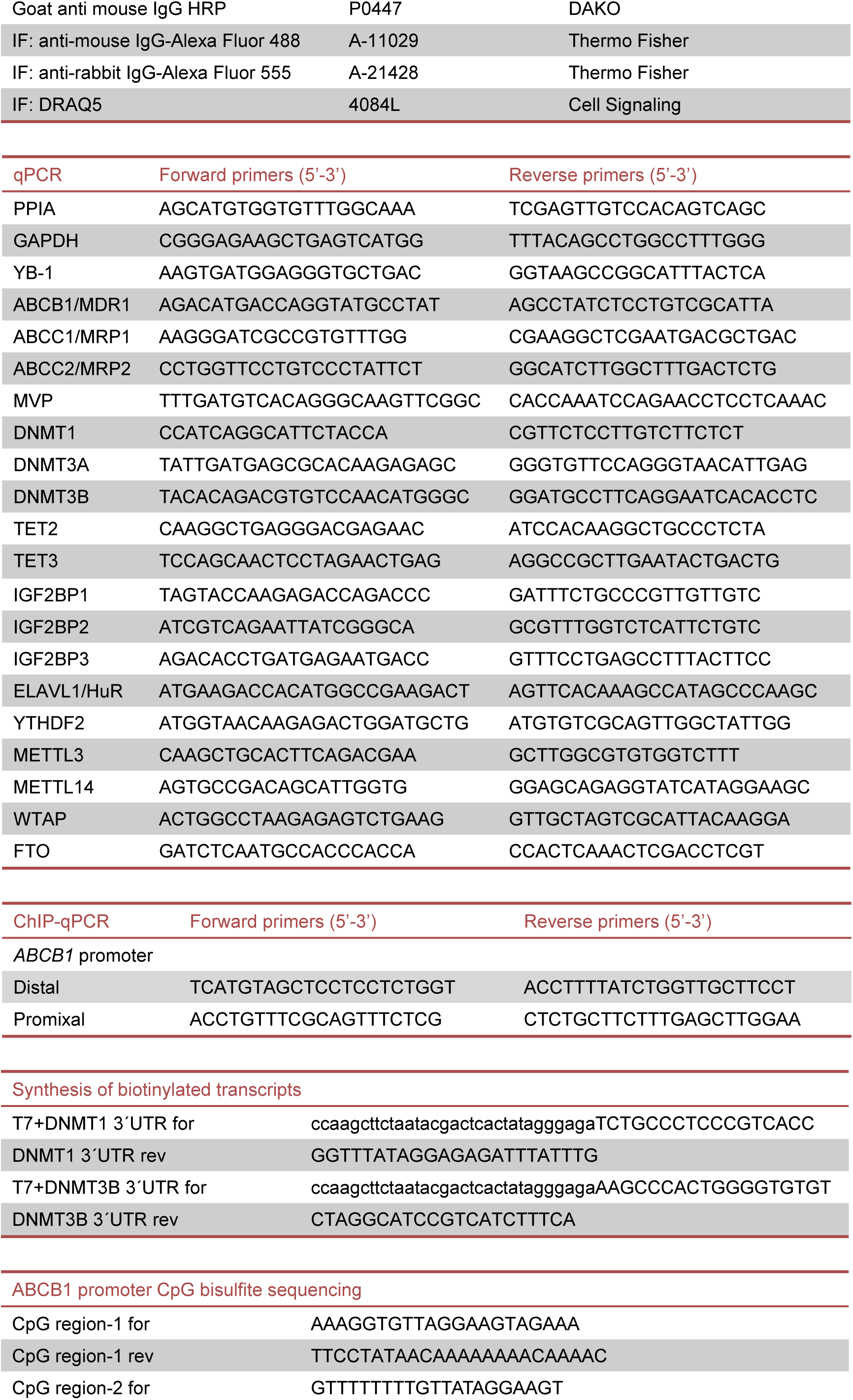
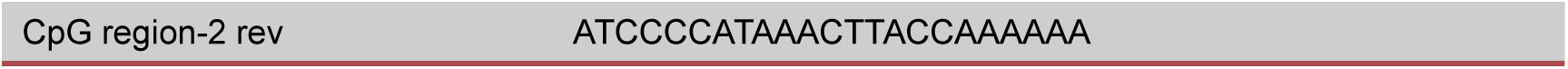

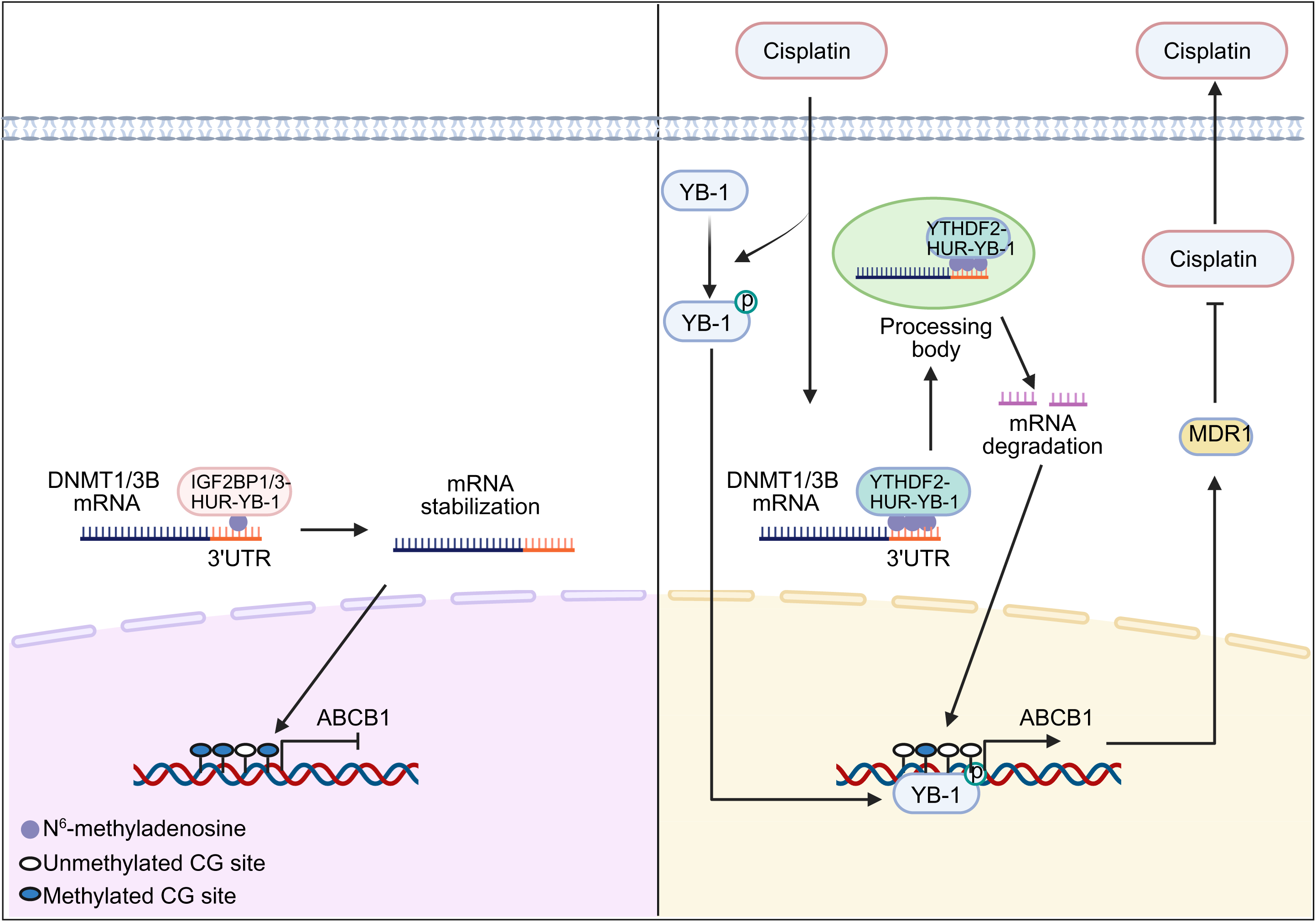

